# Uncovering the ideal plant ideotype for maximising seed yield in *Brassica napus*

**DOI:** 10.1101/2020.12.04.411371

**Authors:** Laura Siles, Kirsty L. Hassall, Cristina Sanchis-Gritsch, Peter J. Eastmond, Smita Kurup

## Abstract

Seed yield is a complex trait for many crop species including oilseed rape (*Brassica napus*), the second most important oilseed crop worldwide. Studies have focused on the contribution of distinct factors in seed yield such as environmental cues, agronomical practices, growth conditions or specific phenotypic traits at the whole plant level, such as number of pods in a plant. However, in spite of the immense economic importance of oilseeds, none of these studies have comprehensively analysed individual traits and their combined contribution to seed yield. Here, we describe the analysis and contribution of 33 phenotypic traits within a *B. napus* diversity set population and their trade-offs on seed yield not only at the whole plant level but also the less studied female reproductive traits. Our results revealed that both winter and spring oilseed rape; the two more economically important oilseed rape groups in terms of oil production; were found to share a common dominant reproductive strategy for seed yield. In this strategy the main inflorescence is the principal source of seed yield, producing a good number of ovules, a large number of long pods with a concomitantly high number of seeds per pod. We observed that winter oilseed rape opted for more reproductive strategies than spring oilseed rape, presenting more environmental flexibility to maximise seed yield. Overall, we conclude that, oilseed rape adopts a similar strategy that is key for maximal seed yield and propose an ideal ideotype highlighting crucial phenotypic traits that could be potential targets for breeding.

**One sentence summary:** The main florescence is the principal source of seed yield in winter and spring oilseed rape, with winter oilseed rape following several reproductive strategies to maximise seed yield.

## Introduction

Improving crop production, particularly seed yield, is vital to ensure food availability for an increasing population in the world. This challenge needs to be met in the face of climate change and reduced availability of arable land. Improving seed yield is a major goal for crop breeding programs for several crop species. *Brassica napus*, also known as rapeseed or oilseed rape (OSR), is the second most important oilseed crop globally (Food and Agriculture Organisation of the United Nations, 2019) accounting for 20% of the world’s total oil production (Hu et al., 2017). It is also a crucial source of high-quality protein for livestock and biofuel production (Raboanatahiry et al., 2018). Therefore, increasing its yield is vital to meet the high demands of oil and animal feed worldwide.

Seed yield in OSR is a complex trait affected by several factors such as environmental cues, agronomical practices, and growth conditions that influence source/sink capacity and resource allocation (Diepenbrock, 2000; Nesi et al., 2008). Studies have focused on the effect of temperature during plant development and growth (Weymann et al., 2015; Brown et al., 2019), plant density and row spacing (Kuai et al., 2015; Ren et al., 2017), nutrient requirements (Stahl et al., 2019), plant and canopy architecture (Bennett et al., 2012; Pinet et al., 2015), pod length (Li et al., 2019) as well as flowering time and petal morphogenesis (Schiessl et al., 2015; Yu et al., 2016) to understand and improve yield in *B. napus*. Given the importance of OSR and complexity of the yield trait, it is surprising that in the last 20 years, studies have focused only on a limited number of phenotypic traits, such as number of pods per plant, number of seed per pod, pod length and number of branches per plant (Habekotté, 1997; Özer et al., 1999; Naazar et al., 2003; Badaran et al., 2007; Tunçtürk and Çiçti, 2007; Başalma, 2008; Sabaghnia et al., 2010; Chen et al., 2014; Ul-Hasan et al., 2014; Moradi et al., 2017; Ahmadzadeh et al., 2019; Tariq et al., 2020). Only one of these studies has focused on 20 phenotypic traits in 49 *B. napus* genotypes (Sabaghnia et al., 2010). Since plant development is complex, any study on seed yield should address the interplay of the various developmental traits and their combined effect.

Seed number per pod (SNPP), pod number and seed weight are considered the most significant components of yield in OSR (Yang et al., 2017), and studies have shed light on the genetic regulation of these traits and their role in seed yield (Li et al., 2015; Yang et al., 2016; Yang et al., 2017; Dong et al., 2018; Li et al., 2019; Zhu et al., 2020). Specifically, SNPP shows a large variation within germplasm resources, from 5 to 35 seeds per pod (Chen et al., 2013). SNPP is determined by the number of ovules per ovary, the proportion of fertile ovules, the number of ovules fertilised and the number of fertilised ovules that develop into seeds (Yang et al., 2016; Yang et al., 2017). However, the natural variation of SNPP and the regulation between ovule number and SNPP in OSR are poorly known, having been explored, so far, only in a limited capacity (Yang et al., 2017). Similarly, there is limited knowledge of the effect, if any, of female reproductive traits, such as ovule number and size and style, ovary and gynoecia length on seed yield (Wang et al., 2011).

Here we present a comprehensive study on the contribution of 33 phenotypic traits and their trade-offs on seed yield, including traits at the whole plant level down to female reproductive traits within a *B. napus* diversity set population formed by 96 genotypes classified in 4 OSR groups subjected to the same vernalisation treatment. We analysed the relationships between the phenotypic traits by Principal Component Analysis (PCA) at the whole population level, performing a Principal Component Regression to relate them to seed yield. Subsequently, a Partial Least Squares (PLS) analysis for Winter OSR (WOSR) and Spring OSR (SOSR), the two more economically important groups of OSR in terms of oil production, was performed. The overall aims of this paper are to study factors influencing seed yield in different OSR groups in a diversity set population and to elucidate the interrelations of these seed yield components. Furthermore, we wanted to identify reproductive strategies that influence seed yield, with a focus on WOSR and SOSR. We unravel the trade-offs between the measured traits at the whole plant level (macrotraits) and in addition, between female reproductive traits (alltraits) and their association to seed production. Finally, we aim to identify the best predictors of seed yield in WOSR and SOSR.

## Results

### Seed yield

Seed yield was measured for the whole diversity set population (Figure 1), presenting values from 3.3 g to 21.3 g per plant. The 4 OSR groups in which the population was divided (see Material and methods section) did not show an even distribution of seed yield (sequential F_3,329_=99.33, *P* < 0.001), with further differences in seed yield observed between lines within each group (F_92,275_=6.01, *P* < 0.001). WOSR and Other groups presented the highest seed yields within the population. The fact that some genotypes within the Other group, presented high seed yield was quite surprising, as these lines are not selected for seed yield, but for their edible leaves or roots. POH 285, Bolko was the highest yielder not only for WOSR, but also for the whole population, meanwhile Tina had the highest seed yield from the Other group. Flash and English Giant were the genotypes with the lowest seed yield for WOSR and Other groups. Mazowiecki and Tapidor DH were the best yielders for SOSR and Semiwinter OSR group, respectively. Meanwhile, Chuanyou 2 and Xiangyou 15 were the genotypes which presented the lowest seed yield not only for Semiwinter OSR group, but for the whole population. Although both WOSR and SOSR genotypes are bred for seed yield, it was observed that, on average, WOSR presented greater seed yield than SOSR (sequential F_1,331_=161.75, *P* < 0.001)(Figure 2), and that SOSR genotypes presented a wider range of seed yield compared to WOSR genotypes, which followed a more symmetric distribution.

**Figure 1.**
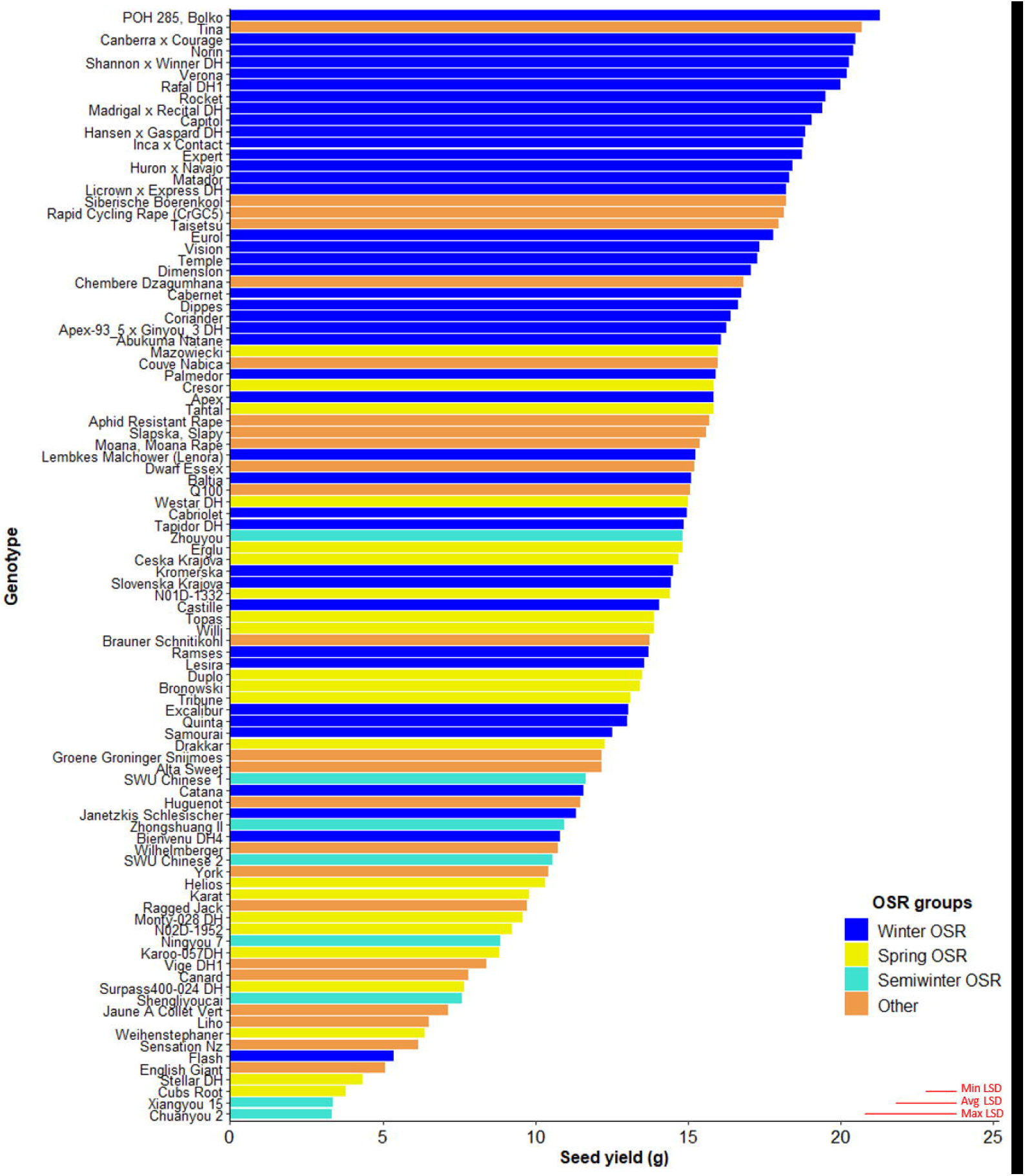
Seed yield (g) for the 96 genotypes of the *Brassica napus* diversity set population for the 4 OSR groups (Winter OSR, Spring OSR, Semiwinter OSR and Other). Data are the mean of 5 biological replicates. Maximum, average and minimum least significant difference (max LSD, avg LSD and min LSD, respectively) are represented as red lines in the bottom right corner of the graph.

**Figure 2:**
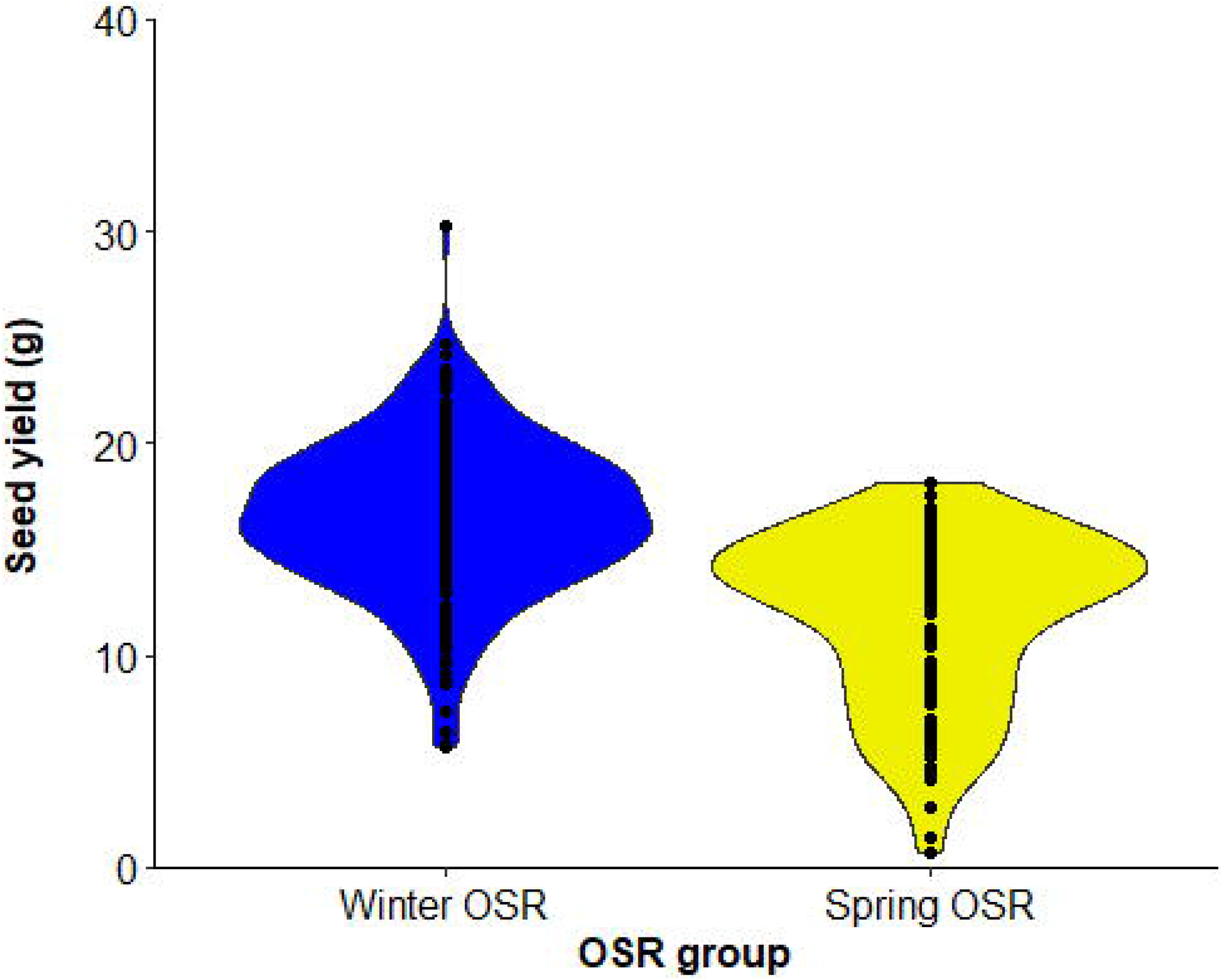
Violin plot for seed yield (g) for Winter OSR and Spring OSR (n=5). Points represent the individual observations for the genotypes in each group.

### Seed yield components

To break down the seed yield trait and determine the interrelation between its components, rank correlations were calculated at macrotrait and alltraits level with a main focus in WOSR and SOSR groups. Pod length was separated into valve and beak length to estimate the contribution of these two phenotypic traits to seed yield. Similarly, gynoecia length was split as ovary and style length. For WOSR_macro_ we found positive correlations between seed yield and seed number (r=0.87) and oil content (r=0.61), with seed number showing the strongest positive correlation with seed yield (Supplemental Figure S1, A). For SOSR_macro_ we found positive correlations between seed yield and seed number (r=0.89), oil content (r=0.85), SNPP_M_ (r=0.70), valve length (r=0.59), pod length (r=0.59), number of pods on a secondary inflorescence (r=0.53) and number of pods in the main inflorescence (r=0.48), and negative correlations between seed yield and thousand grain weight (TGW, r=-0.49), seed area (r=-0.5) and seed area coefficient of variation (r=-0.56) (Supplemental Figure S1, B). SOSR presented higher correlations between seed yield and oil content and SNPP_M_ compared to WOSR. For alltraits, we observed weaker correlations between seed yield and its components (Supplemental Figure S2). For both OSR groups, seed number was the yield component with the strongest correlation with seed yield. We also observed some positive and negative correlations at microtraits level. Hence, the differences in the interrelations between the seed yield components observed in both OSR groups as well as against seed yield suggested different contributions of these phenotypic traits to seed yield.

### Comparison of principal component contribution to seed yield between WOSR and SOSR

The whole diversity set population was included in a PCA as it had a good representation of OSR cultivars that exploit historical recombination between molecular markers and loci associated with trait variation (Harper et al., 2012; Havlickova et al., 2018). This approach enabled us to have an unbiased study at a whole population level. Subsequently, a principal component regression analysis against seed yield was performed to compare the contribution of each principal component (PC) to seed yield for each OSR group as a percentage of total variation explained from all PCs (expressed as contribution to yield (%) herein). Each PC identified combinations of the measured traits explaining the maximal variation in the data, defining ideal reproductive strategies that plants adopt within the population for macrotraits and alltraits, respectively (Supplementary Files S1 and S2). We observed different contribution of PC to seed yield in all groups. As WOSR and SOSR are major seed yielders, we focused our efforts in analysing the differences between these groups. For macrotraits, 12 PCs were identified explaining 95.46% of the variation in the phenotypic traits with associated contribution to seed yield given in Table 1. PC1_macro_ was the reproductive strategy that presented the highest contribution to seed yield in WOSR and SOSR, being the most important reproductive strategy followed by both groups. However, PC1_macro_ contributed ∼1.5 fold more to seed yield in SOSR than in WOSR (78.67% *vs* 54.63%). PC5_macro_ was the next most important reproductive strategy contributing to seed yield for both WOSR and SOSR, but in this case, it explained ∼1.6 fold more contribution to seed yield in WOSR than in SOSR. We observed that PC6_macro_, PC7_macro_ and PC10_macro_ were also contributing to seed yield, albeit more substantially in WOSR compared to SOSR, for which seed yield was largely explained by PC1_macro_ alone. For alltraits, 16 PCs were identified explaining 95.96% of the variation in the phenotypic data with associated contribution to seed yield given in Table 2. Similarly to macrotraits, PC1_alltraits_ was the most important reproductive strategy in both WOSR and SOSR, explaining ∼1.7 fold more contribution to seed yield in SOSR. PC7_alltraits_ and PC6_alltraits_ were the next most relevant reproductive strategies in WOSR and SOSR, presenting a similar contribution to seed yield within each OSR groups but again, explaining more contribution to seed yield in WOSR than in SOSR. For both macrotraits and alltraits, reproductive strategies contributed more to seed yield in WOSR compared to SOSR, for which seed yield was largely explained by PC1_macro_ and PC1_alltraits_.

**Table 1.** Principal Component (PC) contribution to seed yield as a percentage of total variation explained from all PCs for Winter OSR and Spring OSR (expressed as contribution to seed yield (%)) for macrotraits. Percentage is calculated as the ratio of the F-statistic of each PC divided by the sum of F-statistics for all PCs within each OSR group. PCs with small contributions to seed yield are observed.

| Winter OSR |  | Spring OSR |  |
| --- | --- | --- | --- |
| PC<sub>macro</sub> | Contribution to seed yield (%) | PC<sub>macro</sub> | Contribution to seed yield (%) |
| PC1<sub>macro</sub> | 54.63 | PC1<sub>macro</sub> | 78.67 |
| PC2<sub>macro</sub> | 2.96 | PC2<sub>macro</sub> | 0.52 |
| PC3<sub>macro</sub> | 0.02 | PC3<sub>macro</sub> | 0.56 |
| PC4<sub>macro</sub> | 0.91 | PC4<sub>macro</sub> | 0.07 |
| PC5<sub>macro</sub> | 13.11 | PC5<sub>macro</sub> | 8.39 |
| PC6<sub>macro</sub> | 6.80 | PC6<sub>macro</sub> | 6.38 |
| PC7<sub>macro</sub> | 8.46 | PC7<sub>macro</sub> | 2.33 |
| PC8<sub>macro</sub> | 2.48 | PC8<sub>macro</sub> | 0.02 |
| PC9<sub>macro</sub> | 0.43 | PC9<sub>macro</sub> | 0.34 |
| PC10<sub>macro</sub> | 9.16 | PC10<sub>macro</sub> | 2.68 |
| PC11<sub>macro</sub> | 1.01 | PC11<sub>macro</sub> | 0.02 |
| PC12<sub>macro</sub> | 0.05 | PC12<sub>macro</sub> | 0.05 |

**Table 2.** Principal Component (PC) contribution to seed yield as a percentage of total variation explained from all PCs for Winter OSR and Spring OSR (expressed as contribution to yield (%)) for alltraits (macro and microtraits together). Percentage is calculated as the ratio of the F-statistic of each PC divided by the sum of F-statistics for all PCs within each OSR group. PCs with small contributions to seed yield are observed.

| Winter OSR |  | Spring OSR |  |
| --- | --- | --- | --- |
| PC<sub>alltraits</sub> | Contribution to seed yield (%) | PC<sub>alltraits</sub> | Contribution to seed yield (%) |
| PC1<sub>alltraits</sub> | 46.11 | PC1<sub>alltraits</sub> | 76.45 |
| PC2<sub>alltraits</sub> | 1.30 | PC2<sub>alltraits</sub> | 0.81 |
| PC3<sub>alltraits</sub> | 0.68 | PC3<sub>alltraits</sub> | 0.40 |
| PC4<sub>alltraits</sub> | 1.39 | PC4<sub>alltraits</sub> | 1.05 |
| PC5<sub>alltraits</sub> | 0.29 | PC5<sub>alltraits</sub> | 1.13 |
| PC6<sub>alltraits</sub> | 10.06 | PC6<sub>alltraits</sub> | 6.08 |
| PC7<sub>alltraits</sub> | 12.75 | PC7<sub>alltraits</sub> | 6.59 |
| PC8<sub>alltraits</sub> | 1.77 | PC8<sub>alltraits</sub> | 0.02 |
| PC9<sub>alltraits</sub> | 6.05 | PC9<sub>alltraits</sub> | 2.76 |
| PC10<sub>alltraits</sub> | 8.31 | PC10<sub>alltraits</sub> | 1.16 |
| PC11<sub>a1traits</sub> | 1.02 | PC11<sub>a1traits</sub> | 0.03 |
| PC12<sub>alltraits</sub> | 0.44 | PC12<sub>alltraits</sub> | 0.00 |
| PC13<sub>alltraits</sub> | 4.31 | PC13<sub>alltraits</sub> | 0.72 |
| PC14<sub>alltraits</sub> | 3.80 | PC14<sub>alltraits</sub> | 2.19 |
| PC15<sub>alltraits</sub> | 1.69 | PC15<sub>alltraits</sub> | 0.09 |
| PC16<sub>alltraits</sub> | 0.03 | PC16<sub>alltraits</sub> | 0.51 |

### Identification of the most significant reproductive strategies contributing to seed yield within WOSR and SOSR

As described in Tables 1 and 2, there was a total of 12 and 16 PCs for macrotraits and alltraits, respectively, that contribute to seed yield to a larger or smaller extent. To refine this further, a sequential elimination of non-significant terms in the PC regression enabled the identification and order of the most significant reproductive strategies contributing to seed yield within WOSR and SOSR group at macrotraits and alltraits level (Table 3). For macrotraits, WOSR presented 9 PCs, meanwhile SOSR showed 7 PCs that contributed significantly to seed yield. As before, PC1_macro_ was the main reproductive strategy for both WOSR and SOSR, followed by PC5_macro_. For WOSR PC7_macro_ and PC10_macro_ were the next most relevant reproductive strategies contributing to seed yield, whereas PC6_macro_ and PC10_macro_ were the next reproductive strategies for SOSR. For alltraits, WOSR presented 11 reproductive strategies meanwhile we observed 9 for SOSR. While PC1_alltraits_, PC7_alltraits_ and PC6_alltraits_ were the first 3 reproductive strategies for both OSR groups, WOSR presented PC10_alltraits_ while SOSR presented PC14_alltraits_ as important reproductive strategies contributing to seed yield. The higher number of significant PCs by WOSR at both macrotraits and alltraits level confirmed that WOSR presented more reproductive strategies to explain seed yield compared to SOSR. In addition, we observed that the same reproductive strategies present a different order of importance for seed yield between OSR groups.

**Table 3.** Reproductive strategies (PCs) that significantly contribute to seed yield in Winter OSR and Spring OSR for macrotraits and for alltraits (macro and microtraits together) when dropping terms. The order of importance of the reproductive strategies for yield and the approximate F-statistics are reported in the table.

| Macrotraits |  |  |  | All traits |  |  |  |
| --- | --- | --- | --- | --- | --- | --- | --- |
| Winter OSR |  | Spring OSR |  | Winter OSR |  | Spring OSR |  |
| PC order | approximate F-statistics | PC order | approximate F-statistics | PC order | approximate F-statistics | PC order | approximate F-statistics |
| PC1<sub>macro</sub> | 363.61 | PC1<sub>macro</sub> | 413.66 | PC1<sub>alltraits</sub> | 235.2 | PC1<sub>alltraits</sub> | 289.69 |
| PC5<sub>macro</sub> | 117.95 | PC5<sub>macro</sub> | 71.05 | PC7<sub>alltraits</sub> | 68.6 | PC7<sub>alltraits</sub> | 54.71 |
| PC7<sub>macro</sub> | 71.39 | PC6<sub>macro</sub> | 33.59 | PC6<sub>alltraits</sub> | 50.1 | PC6<sub>alltraits</sub> | 35.23 |
| PC10<sub>macro</sub> | 56.01 | PC10<sub>macro</sub> | 16.58 | PC10<sub>alltraits</sub> | 38.14 | PC14<sub>alltraits</sub> | 14.08 |
| PC6<sub>macro</sub> | 44.8 | PC7<sub>macro</sub> | 15.97 | PC14<sub>alltraits</sub> | 29.46 | PC9<sub>alltraits</sub> | 9.98 |
| PC8<sub>macro</sub> | 22.22 | PC3<sub>macro</sub> | 4.77 | PC9<sub>alltraits</sub> | 26.77 | PC5<sub>alltraits</sub> | 8.54 |
| PC2<sub>macro</sub> | 20.16 | PC2<sub>macro</sub> | 1.63 | PC2<sub>alltraits</sub> | 22.49 | PC4<sub>alltraits</sub> | 8.17 |
| PC11<sub>macro</sub> | 8.52 |  |  | PC13<sub>alltraits</sub> | 15.73 | PC13<sub>alltraits</sub> | 6.42 |
| PC4<sub>macro</sub> | 6.38 |  |  | PC15<sub>alltraits</sub> | 9.35 | PC10<sub>alltraits</sub> | 4.94 |
|  |  |  |  | PC8<sub>alltraits</sub> | 8.72 |  |  |
|  |  |  |  | PC4<sub>alltraits</sub> | 7.81 |  |  |

### Reproductive strategies observed in the population for macrotraits and alltraits

Here we present the most important and significant reproductive strategies contributing to seed yield that plants adopt within the diversity set population. We highlighted the combination or trade-offs for the 2 and 3 most important reproductive strategies contributing to seed yield of the measured macrotraits and alltraits, respectively (Tables 4 and 5). Moreover, the other significant PCs contributing to seed yield with a small contribution to seed yield not covered in this section for WOSR and SOSR for macrotraits and alltraits can be found at Supplemental Files S1 and S2, respectively. At the macrotrait level, the main reproductive strategy followed by WOSR and SOSR was PC1_macro_, it being the most important strategy followed by both OSR groups. This reproductive strategy was associated with a reduced number of secondary inflorescences, whereby plants focused their energy and resources mainly in the main inflorescence, and in few secondary branches (Table 4). This strategy was also associated with a high number of pods in the main inflorescence and in secondary inflorescences, presenting a low percentage of pod abortion at the whole plant level. These plants produced long pods in the main inflorescence with a large number of seeds within them. The plants produced a large number of small seeds and with high oil content. Overall, this strategy was associated with high seed yield, with seed number at the whole plant level being the most important trait contributing to seed yield. The next most relevant reproductive strategy (PC5_macro_) was associated with plants producing more flowers in the whole plant, long pods with large uniform circular seeds, but with more seed area coefficient of variation. As in the main reproductive strategy (PC1_macro_), this strategy was associated with high seed oil content. However, in this case, seed area was more important than seed number.

**Table 4.** Winter OSR and Spring OSR reproductive strategies for macrotraits. The traits in the tables are ordered from the most to the least influential trait within the reproductive strategy. The contribution to the PC for each trait is also included, calculated as the percentage of each loading, only showing those traits with an important contribution.

| Reproductive strategy | Positively correlated with seed yield | Negatively correlated with seed yield |
| --- | --- | --- |
| PC1<sub>macro</sub> | SN (6.70%)<br>SNPP<sub>M</sub> (6.48%)<br>OC (5.50%)<br>VL<sub>M</sub> (5.07%)<br>PL<sub>M</sub> (5.01%)<br>SW<sub>M</sub> (4.93%)<br>PN<sub>M</sub> (4.45%)<br>PN<sub>1S</sub> (3.91%) | PA (6.17%)<br>PA<sub>S</sub> (5.92%)<br>PA<sub>1S</sub> (5.90%)<br>PA<sub>M</sub> (5.88%)<br>NI (4.77%)<br>NI-1 (4.66%)<br>FN (4.05%)<br>TGW (3.90%) |
| PC5<sub>macro</sub> | SW<sub>M</sub> (6.39%)<br>SC<sub>M</sub> (6.07%)<br>TGW (5.62%)<br>OC (5.51%)<br>FN (5.39%)<br>SC (4.73%)<br>SA<sub>M</sub> (4.67%)<br>PL<sub>M</sub> (4.52%)<br>VL<sub>M</sub> (4.42%)<br>SA (4.09%) | SAvar (7.55%) |

**Table 5.** Winter OSR and Spring OSR reproductive strategies for all traits (macro and microtraits together). The traits in the tables are ordered from the most to the least influential trait within the reproductive strategy. The contribution to PC for each trait is also included, calculated as the percentage of each loading. In this case, traits with lower contributions were also included in order to elucidate relationships with microtraits.

| Reproductive strategy | Positively correlated with seed yield | Negatively correlated with seed yield |
| --- | --- | --- |
| PC1<sub>alltraits</sub> | SNPP<sub>M</sub> (5.92%)<br>SN (5.82%)<br>VL<sub>M</sub> (4.84%)<br>PL<sub>M</sub> (4.82%)<br>OC (4.66%)<br>SW<sub>M</sub> (4.63%)<br>PN<sub>M</sub> (4.11%)<br>PN<sub>1s</sub> (3.53%)<br>BL (3.46%)<br>PN (2.37%)<br>ON (1.80%) | PA (5.70%)<br>PA<sub>M</sub> (5.64%)<br>PAs (5.54%)<br>PA1s (5.46%)<br>NI (4.27%)<br>NI-1 (4.14%)<br>FN (3.88%)<br>TGW (3.20%) |
| PC7<sub>alltraits</sub> | ON (7.90%)<br>SN (6.84%)<br>OL (6.83%)<br>GL (6.67%)<br>OC (5.90%)<br>PA (3.97%)<br>PA<sub>s</sub> (3.89%)<br>FN (3.76%) | BL (4.91%)<br>SC (4.18%) |
| PC6<sub>alltraits</sub> | TGW (6.08%)<br>SW<sub>M</sub> (5.38%)<br>SA<sub>M</sub> (5.33%)<br>SA (4.82%)<br>FN (4.50%)<br>PL<sub>M</sub> (4.39%) | SAvar (5.16%)<br>OL (4.00%)<br>TF (3.45%)<br>OAvar (3.26%)<br>GL (3.22%)<br>ON (2.35%) |
|  | VL<sub>M</sub> (4.16%)<br>BL (3.88%)<br>OC (3.85%) |  |

The analysis was extended to include microtraits to assess whether these traits significantly influenced the macrotraits and or seed yield (Table 5). The main reproductive strategy for both WOSR and SOSR including all traits (PC1_alltraits_) was similar to PC1_macro_. Moreover, this strategy was associated with plants presenting long beaks and a high number of ovules in the main inflorescence. The next reproductive strategy (PC7_alltraits_) was associated with plants with short beaks but with high number of ovules, long ovaries and long gynoecia, with these traits presenting a high contribution within the reproductive strategy. These plants produced a high number of flowers and displayed pod abortion. This strategy was associated with plants generating a large number of seeds with high seed oil content. Finally, the next most relevant reproductive strategy (PC6_alltraits_) was similar to PC5_macro,_ with the addition of being associated with short ovaries and gynoecia, low number of ovules, long beaks and seeds with high oil content.

### High yielders follow several reproductive strategies

WOSR and SOSR genotypes were ranked for each reproductive strategy for macrotraits and alltraits in order to identify whether consistently high yielding OSR follow a certain strategy. WOSR genotypes POH 285, Bolko; Canberra x Courage, Norin, Shannon x Winner DH and Verona and Spring OSR genotypes Mazowiecki, Cresor, Tantal, Westar DH and Erglu were identified as high yielders. We observed that high yielders in both OSR groups did not follow a particular reproductive strategy for macrotraits or alltraits, but a combination of them, as suggested by our results (Supplemental Tables S1, S2, S3 and S4). However, consistent with our analyses, they all presented a good rank for PC1_macro_ and PC1_alltraits_.

Interestingly, the five worst WOSR yielders Flash, Bienvenu DH4, Catana, Samourai and Quinta showed low adoption of the main reproductive strategy PC1 in both macrotraits and alltraits level. For SOSR, as observed in WOSR, the worst five worst yielders, Cubs Root, Stellar DH, Wiehenstephaner, Surpass400-024DH and Karoo-057-DH also presented a low rank for PC1_macro_ and PC1_alltraits_.

### A PLS analysis corroborates the main strategy for WOSR and SOSR, and seed number is the best predictor of seed yield

Our initial analyses at the whole population level highlighted a distinctive response between WOSR and SOSR in terms of reproductive strategies relevant to seed yield. Subsequently, WOSR and SOSR groups were analysed separately to fully capture the strategies employed by each. A PLS analysis for WOSR and SOSR was performed in order to corroborate the results obtained at whole population level and to determine the best predictor of seed yield for WOSR and SOSR, respectively. The PLS approach iteratively identifies combinations of traits, defining the PLS components that are maximally related to seed yield and then combines these components to get an overall assessment of the contribution of each trait to seed yield. For the macrotraits, 9 components explained 96.3% and 97.3% of the variation in seed yield in WOSR and SOSR, respectively (Supplemental Table S5). We observed that although both OSR groups presented the same number of components (chosen by cross-validation), the contribution to seed yield from component 1 was substantially higher in SOSR, explaining 74% of the variation in seed yield. Component 1 presented the same combination of significant traits for PC1_macro_ and PC1_alltraits_, confirming that this was the main reproductive strategy contributing to high seed yield at both macrotraits and alltraits level. Component 1 also presented the highest variation in seed yield for WOSR (44.2%), but other components were also represented to a large extent, supporting the idea that WOSR adopt more reproductive strategies for optimising seed yield than SOSR. The same trends and results were observed for alltraits. For WOSR, 11 components explained 97.0% of the variation on seed yield, while 8 components explained 96.8% of the variation in seed yield for SOSR (Supplemental Table S6).

Taking account of all the components contributing to seed yield, the most important trait affecting seed yield in WOSR and SOSR for macrotraits and WOSR alltraits was seed number, followed by TGW, both positively associated with yield. On the other hand, the predictors most negatively associated with seed yield were number of flowers, number of pods on secondary inflorescences and number of secondary inflorescences in WOSR. Whereas for SOSR they were time to flowering, pod abortion in the whole plant and seed compactness from 10 pods from the main inflorescence (Supplemental Table S7).

### The number of seeds per pod increases as valve lengthens

As seed number was the best predictor of seed yield, and SNPP and pod length presented a high contribution in the main reproductive strategy followed by WOSR and SOSR, we investigated whether the number of seeds increased as the pods valves lengthen (Figure 3). We observed that the SNPP increased as valve length increased following a similar pattern in WOSR and SOSR, presenting an exponential increase until approximately 5 cm of valve length, followed by a more linear increase. We observed the same trend for all SOSR genotypes with one exception, Karat. Interestingly, Semiwinter OSR genotypes presented, in general, long valves with fewer seeds, which was especially evident in Xiangyou 15 and Zhongshuang II. This highlights the fact that the selection of long pods needs to be linked to good seed packing. On the other hand, we observed some WOSR genotypes, such as Kromerska and Hansen x Gaspard DH, that presented shorter valves with a high SNPP. Interestingly, Hansen x Gaspard DH also presented long valves with a low number of seed, and in particular this genotype exhibited a high variability in SNPP. The SNPP coefficient of variation presented a wider distribution for SOSR compared to WOSR (Figure 4A), but on average, both groups presented no significant differences for this trait (sequential F_1.339_=0.92, *P* = 0.337), demonstrating that this trait is as variable in both OSR groups. In general, WOSR genotypes presented bigger seed areas than SOSR (sequential F_1.318_=151.84, *P* < 0.001, Figure 4B), presenting a maximum around 3.2 mm^2^ with a skewness towards bigger seeds. However, SOSR genotypes seemed to produce two types of seeds, one around 2.7 mm^2^ and other around 3.5 mm^2^, presenting a multimodal distribution. Finally, SOSR produced less uniform seed areas compared to WOSR (sequential F_1.313_=21.02, *P* < 0.001, Figure 4C).

**Figure 3:**
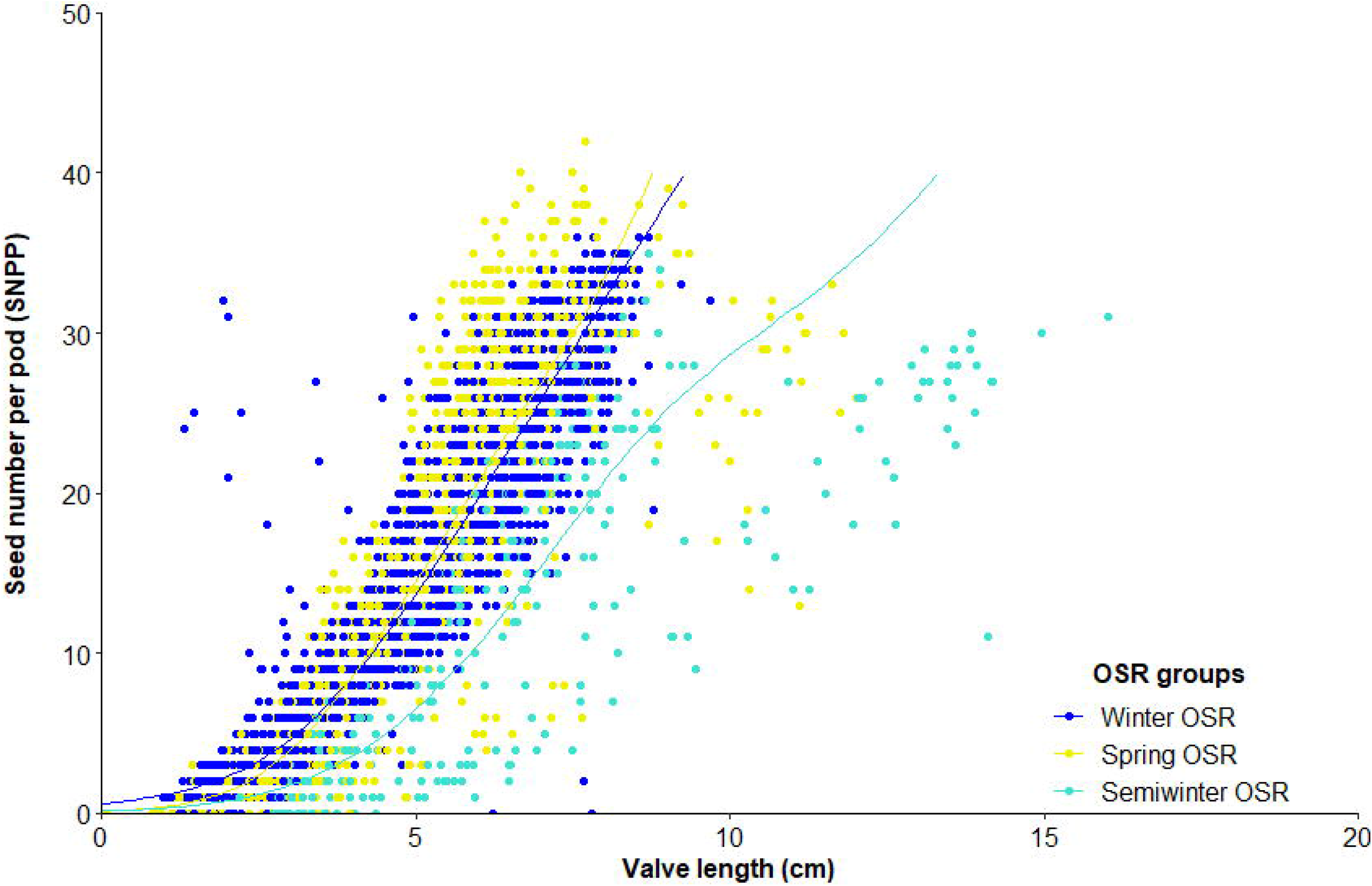
Relationship between seed number/ pod (SNPP) and valve length from 10 pods from the main inflorescence for Winter OSR, Spring OSR and Semiwinter OSR. Fitted lines are the result of a generalized additive mixed model.

**Figure 4:**
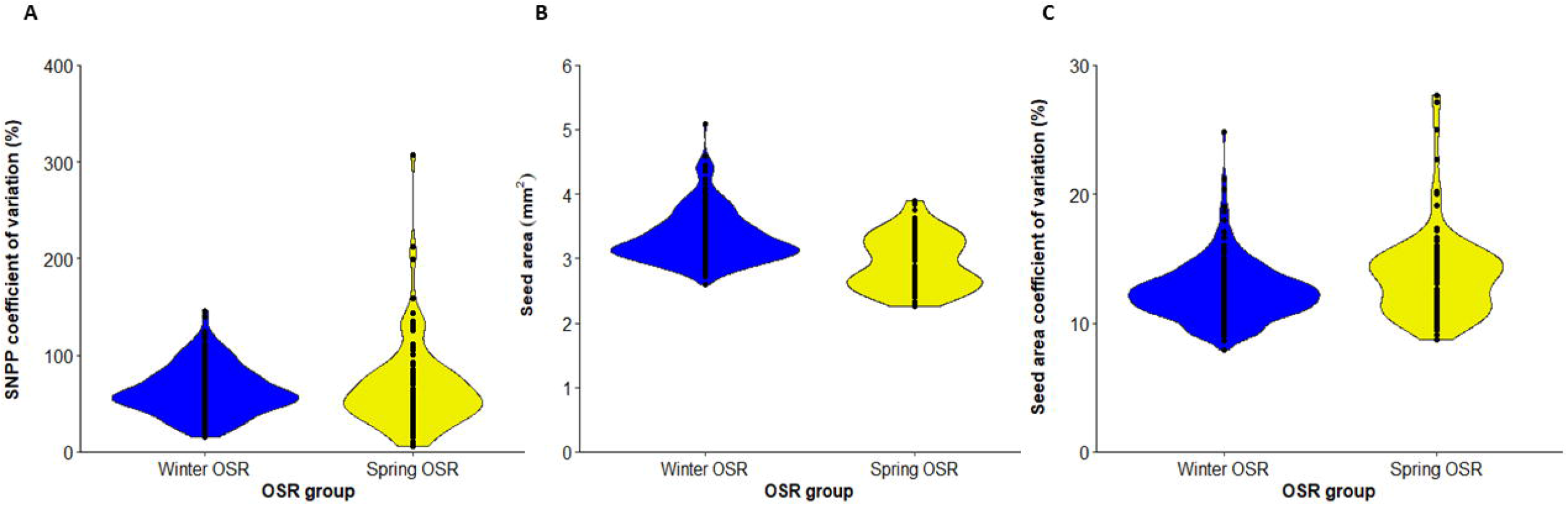
Violin plots for A) seed number/ pod (SNPP) coefficient for variation (%), B) seed area (mm^2^) and C) seed area coefficient of variation (%) for Winter OSR and Spring OSR (n=5). Points represent the individual observations for the genotypes in each group.

## Discussion

The differences in seed yield observed for the OSR groups in the diversity set population can be explained by varying combinations of reproductive strategies adopted by these groups. Our analyses highlighted distinct differences in the contribution to seed yield arising from different reproductive strategies, with PC1_macro_ and PC1_alltraits_ providing the biggest contribution to seed yield, especially evident in SOSR. The seeds from the main inflorescence were the principal source of seed yield for WOSR and SOSR. This strategy was associated with a reduced number of secondary inflorescences, presumably with the plants relocating their carbon assimilates primarily to the main inflorescence. The above result highlights the importance that plant architecture may play in assimilate partitioning among plant organs. The successful development of pods and seeds and their variation in number is determined by the quantity of assimilates available at the whole plant level and the competition with other developing organs (Arathi et al., 1999; Diepenbrock, 2000). This is particularly crucial during the plant reproductive phase, when competition between developing pods and seeds among different inflorescences occurs, causing a high demand of carbon assimilates within a short period of time (Wang et al., 2011). Consequently, the reduction of number of flowering inflorescences decreases intra-plant competition that may be responsible for loss of buds, flowers and seeds (Diepenbrock, 2000), resulting in a high number of pods in the main inflorescence with reduced percentage of pod abortion and enhanced seed yield. Leaves are the major source of photosynthesis in OSR until flowering, providing assimilate source supporting pod growth. At the onset of flowering, leaf area decreases due to canopy shading and flower photon reflectivity and leaves start to fall, reducing leaf photosynthesis by 40% (Diepenbrock, 2000). Therefore, long pods enhance photosynthetic capacity as the developing pod wall become the main intercept of solar radiation, contributing up to 70% of the assimilates to seed filling (Diepenbrock, 2000; Li et al., 2019). This is in concordance with our results, in which we observed that longer valves can hold a higher number of seeds, and that a high number of pods with long valves with a high SNPP were associated with high seed yield. Previous studies have also found that number of pods per plant and SNPP in *Brassica* sp. genotypes were major contributors to seed yield (Özer et al., 1999; Badaran et al., 2007; Tunçtürk and Çiçti, 2007; Chen et al., 2014; Ul-Hasan et al., 2014; Moradi et al., 2017; Ahmadzadeh et al., 2019; Tariq et al., 2020). Our study further confirms that seed number is the single most important trait affecting seed yield. Specifically, Başalma (2008) also reported that the number of pods in the main inflorescence rather than the whole plant presented a positive correlation with seed yield in WOSR. Within the main reproductive strategy that WOSR and SOSR were following, we observed a trade-off between seed number and seed size, as the plants produce high number of seeds at the expense of seed size. This can again be explained by resource availability in the mother plant, with plasticity in seed number proving more beneficial in an environment of variable resource availability (Sadras, 2007).

Interestingly, when the microtraits were included in the analyses, we observed that long beaks and a high number of ovules were also associated with the main reproductive strategy, highlighting the importance of these often-ignored phenotypic traits. A high number of ovules is essential to obtain a final high number of seeds, the trait affecting seed yield maximally in the main reproductive strategy. Although PC1 was the main reproductive strategy for both OSR groups, other reproductive strategies presented significant contribution to seed yield albeit to a lesser extent. These strategies highlighted the importance of the main inflorescence by producing long pods with big seeds at the macrotraits level. When the microtraits were included, we observed the importance of producing a high number of ovules with long ovaries and gynoecia at expense of beak length and seed compactness for one strategy (PC7_alltraits_) or generating long pods with big seeds with short ovary and gynoecia lengths. Although these reproductive strategies presented less contribution to seed yield than PC1, the fact that WOSR retained more of these strategies compared to SOSR in both macrotraits and alltraits level was an important difference between these two OSR groups, which can be associated to their different life cycles. WOSR requires vernalisation to promote the onset of flowering, being grown largely in Western Europe and United Kingdom, where winters are mild. Their seeds are sown in later summer and survive winter in a leaf rosette form, putting a lot of effort in vegetative growth. They flower between March and May, completing the development of pod and seeds by the end of June (Diepenbrock, 2000; Nesi et al., 2008; Brown et al., 2019). On the other hand, SOSR genotypes present a faster life cycle and are cultivated in Canada, Australia, Asia and Eastern Europe. In these countries, winters are too cold and SOSR genotypes are sown at the end of winter as they are not vernalisation dependent (Snowdon et al., 2007; Nesi et al., 2008). The differences in the life cycle and temperatures the plants are subject to appear to be the main drive of varying reproductive strategies, as WOSR cultivars experience more variable environmental conditions during their life cycle. Moreover, as its life cycle is longer than the SOSR, they have more time to adapt and compensate for environmental or mechanical damages; as for example frost events at the onset of flowering (Lardon and Triboi-Blondel, 1995), periods of high temperatures during flowering that can cause a reduction in pollen viability and germinability and pod abortion (Angadi et al., 2000; Young et al, 2004) or water stress during flowering (Champolivier and Merrien, 1995; Elferjani and Soolanayakanahally, 2018); and hence secure reproductive success. The plasticity presented by WOSR may explain why WOSR genotypes have higher seed yields than SOSR.

The PLS analysis corroborated that the main inflorescence was the main contributor to seed yield in both OSR groups (PC1), and that although PC1 was the single larger reproductive strategy in WOSR contributing to seed yield, and among the studied genotypes, WOSR presented more reproductive strategies in order to explain seed yield. The most important traits contributing to seed yield in PLS components 2 and 3 after having accounted for the association between the components and seed yield, highlighted common traits with the significant reproductive strategies contributing to seed yield. Furthermore, seed number was the best predictor of seed yield for WOSR and SOSR, followed by TGW as a proxy of seed size, confirming the results observed at the whole population level.

Here we propose that an ideal SOSR or WOSR phenotype (ideotype) for high seed yield should have a limited number of inflorescences with a good number of ovules and pods on the main inflorescence and reduced percentage of pod abortion. The pods should have long valves with high SNPP for producing seeds with high seed oil content (Figure 5A). The WOSR ideotype can also invest in more flowers, a few secondary inflorescences and bigger seeds in pods with long valves to produce high seed yields (Figure 5B). Both SNPP and pod length are phenotypic traits that present relatively high heritability, therefore are important targets for breeding selection (Shi et al., 2009; Zhang et al., 2011; Li et al., 2019) as they still present great variation in OSR germplasms resources. However, it is important to highlight that long pods in itself are not sufficient but should demonstrate good seed packing for maximal seed yield. It remains to be determined whether SNPP is subject to genetic control independent of ovule number.

**Figure 5:**
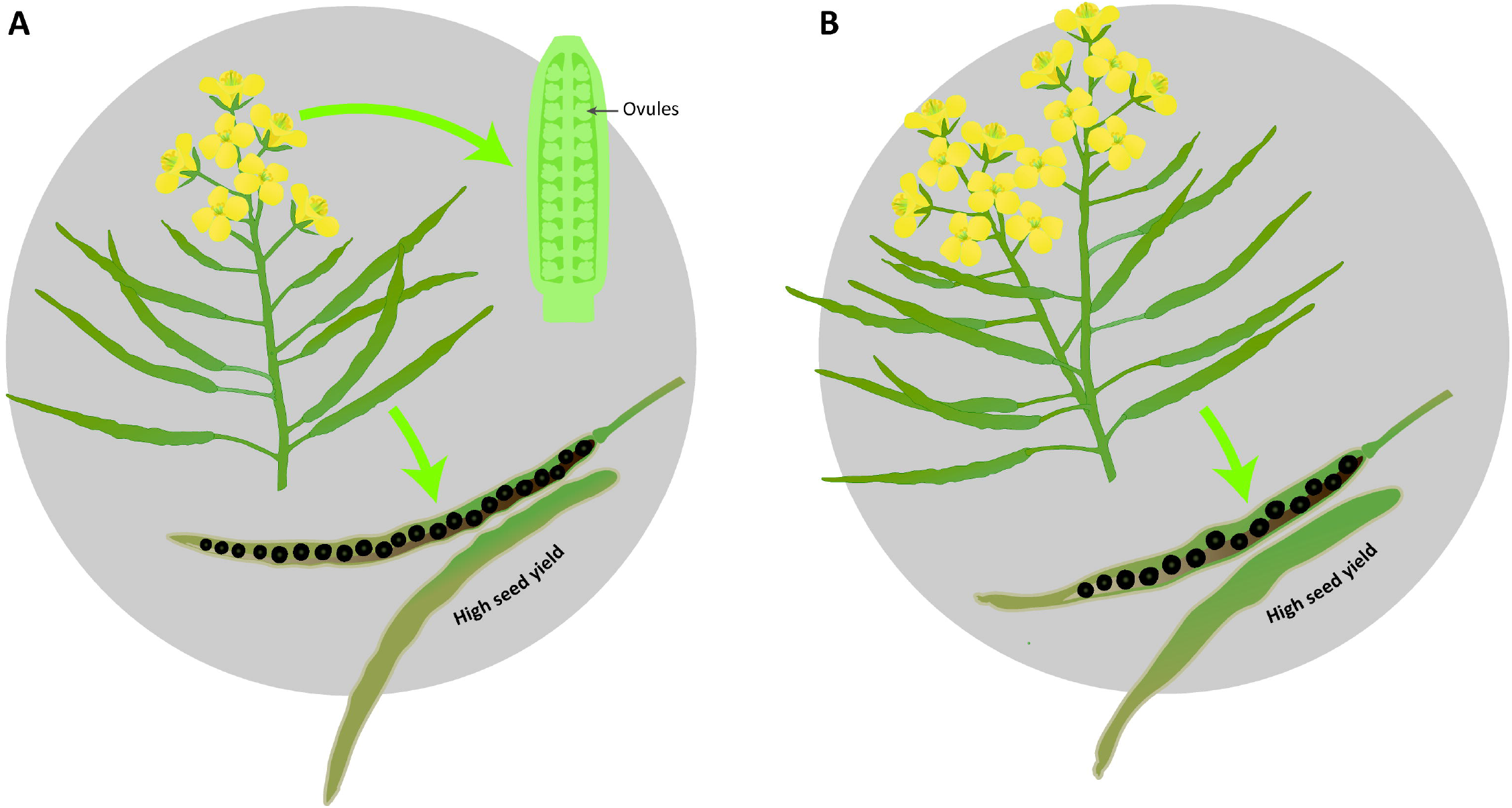
Graphical representation of the proposed ideotypes of *Brassica napus* for obtaining maximal seed yield. A) Ideal ideotype for SOSR and WOSR. B) Additional WOSR ideotype leading to high seed yield.

## Conclusions

Our study uncovered that in spite of the genetic diversity represented across *Brassica* sp. genotypes, OSR follow primarily one discrete strategy for maximal seed yield. We examined different reproductive strategies followed by WOSR and SOSR groups in order to achieve high seed yields from the whole plant level down to female reproductive traits. WOSR can follow different reproductive strategies to maximise its yield although PC1 is the predominant strategy contributing to seed yield in this OSR group. Although OSR plants demonstrate large differences in vernalisation, branching, flowering time and canopy structure, they appear to uniformly prefer a single approach for seed yield. This knowledge is important for breeders in determining target traits for improvement that can confer maximum yield benefit in OSR.

## Material and methods

### Plant material and growth conditions

The *B. napus* diversity set population consisted of 96 genotypes that included WOSR, SOSR, Semiwinter OSR, swede, kales, unspecified and Spring and Winter fodder genotypes (Harper et al., 2012; Havlickova et al., 2018). The population was classified in 4 OSR groups, including WOSR (42 lines), SOSR (22 lines), Semiwinter OSR (8 lines) and Others (24 lines which included swede, kale, unspecified and fodder genotypes, Supplemental Table S8). The seeds were germinated in P24 trays with John Innes Cereal Mix as described in (Siles et al., 2020).When the plants presented 4 true leaves, they were transferred to a vernalisation room with an 8h photoperiod at 4°C day/night for 8 weeks. The plants were re-potted in 2L pots with John Innes Cereal Mix and were allocated in two glasshouse compartments in long-day conditions (16 h photoperiod) at 18°C day/ 15°C night (600w SON-T, high pressure sodium lighting). Plants were grown on ebb and flow benches, flood watered twice a day for 25 minutes. Once the plants started to mature, watering was reduced to once a day, decreasing the time of watering gradually until turning the water off completely. Perforated bread bags (380 mm x 900 mm, WR Wright & Sons Ltd, Liverpool, UK) were used to enclose inflorescences to prevent cross-pollination from neighbouring plants once the plants started to bolt.

### Phenotyping

A total of 33 traits and seed yield were measured for the entire diversity set population, performing a total of 14,976 measurements. Seed yield and a further 26 phenotypic traits, measured on all 5 biological replicates of each genotype, were classified as macrotraits as they could be measured at the whole plant level. The other 7 phenotypic traits were classified as microtraits, as these required some level of dissection being measured, performing 3 biological replicates for each genotype. The combination of macrotraits and microtraits were classified as alltraits. A list of the names, units and abbreviations used for the 33 measured phenotypic traits and seed yield can be found in Supplemental Table S9.

### Macrotrait phenotyping

Plants were monitored daily visually, and time to flowering was recorded. Time to maturity, plant height, number of flowering and secondary inflorescences, number of pods and percentage of pod abortion in the main inflorescence were manually measured and counted. Based on 2 representative secondary inflorescences, the number of pods and the percentage of aborted pods for a single secondary inflorescence were determined. Moreover, we estimated the number of pods and percentage of aborted pods for all secondary inflorescences. The number of flowers on the whole plant was estimated by the number of pods on the whole plant.

Ten consecutive pods per plant from the main inflorescence between the 9^th^ and the 19^th^ pod were imaged (Nikon D5300, HOYA Pro1 Digital 52mm MC UV objective). Subsequently, each pod was opened to remove the seeds, which were placed in individual petri dishes in order, and imaged. Pod and valve length were measured using SmartRoot tool in Image J 1.48v, and their average was calculated for each plant. The number of seeds per pod (SNPP_M_) was counted using Cell counter tool in Image J, and its average was calculated for each plant. Seed area and compactness (a measure of the circularity of the seed) from seeds from 10 pods from the main inflorescence and from the whole plant were recorded (Videometer, Videometer A/S, Herlev, Denmark). For each plant, 3 technical reps were measured, and seed area and compactness were averaged for each plant.

Seed oil content was measured by time-domain nuclear-magnetic resonance (TD-NMR, Bruker minispec mq-20 NMR, Bruker, Massachusetts, USA) for each plant (standardised by seed moisture content at 9%). TGW was calculated from a sample of 200 seeds from each plant, and the number of total seeds per plant was estimated by TGW. Finally, seed weight from 10 pods from the main inflorescence as well as from the whole plant (seed yield) were obtained.

### Microtrait phenotyping

a total of 3 buds per plant at stages 12-13 (Sanders et al., 1999) were collected 24 hours prior to anthesis (pre-fertilization stage) between buds 6 and 20 from the main inflorescence for 3 biological reps per genotype. Sepals, petals and anthers were removed, obtaining 3 gynoecia per plant placed in a glass vial with 4% paraformaldehyde in 0.01M Phosphate Buffer Saline and stored at 4°C until further processing. For each plant, an image of the 3 gynoecia using a stereo microscope (Leica M-205, Leica microsystems) was captured. Then, the ovules were extracted from the ovaries and imaged. Ovary, style and gynoecia length as well as ovule area and number were measured from these images using Image J. For each plant, the average of 3 technical reps was measured. Beak length from 10 pods from the main inflorescence was measured using SmartRoot tool in Image J, and its average was calculated for each plant.

### Ovule, seed area and seed number per pod coefficient of variation

Each biological replicate contained between 70 and 120 ovule measurements taken from 3 gynoecia (around 30-40 measurements per gynoecia). Consequently, the percentage coefficient of variation of ovule area was calculated for each plant,

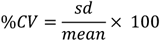

where sd is the standard deviation of all ovule measurements (within a single plant) and mean is the average ovule area. Similarly, the percentage coefficient of variation of seed area was calculated per plant, where between 300-1200 measurements were available per plant, and the coefficient of variation for SNPP was calculated from 10 pods per plant with a small number of exceptions (1 plant had 6 pods and 2 plants had 9 pods).

### Statistical analyses

#### Statistical Design

96 genotypes with 5 biological replicates were arranged in 2 glasshouses each according to a non-resolvable row-column design.

#### Univariate Analysis

Each trait was analysed using a linear mixed model. The block structure was defined by glasshouse/(row x column), and the main effect of glasshouse was fitted as a fixed effect. Glasshouse.row and glasshouse.col were both fitted as random effects. The treatment term accounting for differences between genotypes was fitted as a fixed effect, with statistical significance assessed by the Kenward-Roger approximate F-tests (Kenward and Roger, 1997) after having fitted the main effect of glasshouse. Further refinement of the random model was done on a trait-by-trait basis, and where necessary, variables were transformed to satisfy homogeneity of variance (Supplemental Table S10).

The three percentage abortion traits (main inflorescence, secondary inflorescences and whole plant) were analysed on the logit scale with the associated number of pods (on main inflorescence, on secondary inflorescences and on whole plant, respectively) included as a weight. For the 23 macrotraits (all 26 excluding the 3 weighted abortion traits) independent AR(1)-AR(1) correlated error structures were imposed on the rows and columns of each glasshouse.

#### Principal Component Analysis (PCA)

PCA was performed on i) the set of 26 macrotraits (PCA_macro_) and ii) the set of 33 microtraits (PCA_alltraits_) using the NIPALS algorithm implemented in the mixOmics package of R (Rohart et al., 2017) and run using the correlation matrix. Input variables were adjusted for glasshouse and position within glasshouse as per the univariate analysis and kept on the transformed scale where applicable. For PCA_macro_, 12 principal components (PCs) were retained, explaining 95.46% of the variation in the data. For PCA_alltraits_, 16 PCs were retained, explaining 95.96% of the variation in the data.

#### Principal component regression

To understand which traits were associated with the observed yield differential (the variation in seed yield), a principal component regression analysis was carried out. This consisted of two parts i) for the macrotraits only, using PCA_macro_ and ii) for alltraits subsetting the data to 3 replicates per genotype using PCA_alltraits_. For the macrotraits, a baseline model for seed yield was defined as per the above univariate analysis. Specifically, a linear mixed model with random model defined by glasshouse.(row x column) and fixed model defined by glasshouse + genotype. Two additional auto-correlated error terms were fitted across the rows and across the columns within each glasshouse to further account for the spatial dependence. The principal component regression models kept the same random structure with correlated error terms, but with fixed model consisting of glasshouse + OSRgroup * (PC1 + PC2 + … + PC12). Significance of individual terms was assessed by the marginal Kenward-Roger F-statistic (Kenward and Roger, 1997). An approximate percentage variance each model accounted for was calculated according to,

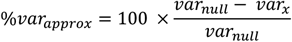

where *var*_*null*_ is the sum of the variance components under a model with no fixed effects beyond Glasshouse and *var*_*x*_ is the sum of the variance components under a model with a defined fixed model (Welham et al, 2015). For the combined set of macro and microtraits, restricted to the 3 replicates per genotype, the principal component regression modelling was performed in the same way as above, with the exception that no autocorrelated spatial error terms were included in the mixed models and a maximum of 16 PCs were allowed. Analysis of the contribution of each PC to seed yield was compared across OSR groups by the associated Kenward-Roger F-statistic. Specifically, for each PC regression model, the F-statistics of the saturated model were expressed as a percentage of the sum of all F-statistics for the PCs within each OSR group. To identify the minimal set of important PCs for determining seed yield, the above PC regression models were refined through a sequential backwards elimination process removing any term found to be non-significant (at a 5% threshold).

#### Partial Least Squares (PLS)

PLS regression models were fitted to the subsets of WOSR and SOSR genotypes separately. Analyses were performed on all macrotraits (173 and 100 observations for WOSR and SOSR, respectively) and on alltraits (106 and 60 observations for WOSR and SOSR, respectively). Both the response (seed yield) and explanatory variables were standardised (mean centred and scaled by the standard deviation) and the PLS2 algorithm was used. Only observations with a complete set of measured traits were included.

#### Modelling seed number per pod

A model was fitted to the SNPP to explore the relationship between SNPP and valve length. Generalized additive mixed models were fitted to the data using the gamm4 package in R. Random effects of glasshouse/(row*col) were included and a separate thin-plate regression spline was fitted to each OSR type.

Linear mixed models (both univariate and PC regressions) and Partial least squares analysis was done using Genstat 20^th^ Edition. Principal components analysis and generalized additive mixed models were done using R statistical software environment v3.6.1.

## Supplemental Material

Supplemental Figure S1: Spearman’s correlation for macrotraits for A) Winter OSR and B) Spring OSR groups (n=5). PH=plant height (cm), NI= number of flowering inflorescences, NI-1= number of secondary inflorescences, TF= time to flowering (days), FN= number of flowers on the whole plant, PN_M_=number of pods on the main inflorescence, PN_1S_=number of pods on a secondary inflorescence, PN_S_=number of pods on secondary inflorescences, PN= number of pods on the whole plant, PA_M_= pod abortion on the main inflorescence (%), PA_1S_=pod abortion on a secondary inflorescence (%), PA_S_=pod abortion in secondary inflorescences (%), PA=pod abortion in the whole plant (%), TM= time to maturity (days), PL_M_=pod length from 10 pods from the main inflorescence (cm), VL_M_= valve length from 10 pods from the main inflorescence (cm), SNPP_M_= seed number/ pod from 10 pods from the main inflorescence, SA_M_= seed area from 10 pods from the main inflorescence (mm^2^), SC_M_= seed compactness from 10 pods from the main inflorescence, SW_M_= seed weight from 10 pods from the main inflorescence (g), SA= seed area from the whole plant (mm^2^), SC= seed compactness from the whole plant, SAcvar= seed area coefficient of variation from whole plant (%), TGW= thousand grain weight (g), SN= estimated total seed number from the whole plant (by TGW), OC= seed oil content from the whole plant (%), SY= seed weight from the whole plant (seed yield, g).

Supplemental Figure S2: Spearman’s correlation for alltraits for A) Winter OSR and B) Spring OSR groups (n=3). PH=plant height (cm), NI= number of flowering inflorescences, NI-1= number of secondary inflorescences, TF= time to flowering (days), FN= number of flowers on the whole plant, ON=ovule number, OA=ovule area (mm^2^), OAcvar=ovule area coefficient of variation (%), OL= ovary length (mm), GL=gynoecia length (mm), SL= style length (mm), PN_M_=number of pods on the main inflorescence, PN_1S_=number of pods on a secondary inflorescence, PN_S_=number of pods on secondary inflorescences, PN= number of pods on the whole plant, PA_M_= pod abortion on the main inflorescence (%), PA_1S_=pod abortion on a secondary inflorescence (%), PA_S_=pod abortion in secondary inflorescences (%), PA=pod abortion in the whole plant (%), TM= time to maturity (days), PL_M_=pod length from 10 pods from the main inflorescence (cm), VL_M_= valve length from 10 pods from the main inflorescence (cm), BL= beak length (cm), SNPP_M_= seed number/ pod from 10 pods from the main inflorescence, SA_M_= seed area from 10 pods from the main inflorescence (mm^2^), SC_M_= seed compactness from 10 pods from the main inflorescence, SW_M_= seed weight from 10 pods from the main inflorescence (g), SA= seed area from the whole plant (mm^2^), SC= seed compactness from the whole plant, SAcvar= seed areacoefficient of variaiton from whole plant (%), TGW= thousand grain weight (g), SN= estimated total seed number from the whole plant (by TGW), OC= seed oil content from the whole plant (%), SY= seed weight from the whole plant (seed yield, g).

Supplemental File S1. Principal Component (PC) loadings for the significant macrotraits reproductive strategies retained by Winter OSR and Spring OSR groups.

Supplemental File S2. Principal Component (PC) loadings for the significant alltraits reproductive strategies retained by Winter OSR and Spring OSR groups.

Supplemental Table S1: Ranks the Winter OSR genotypes according to its position within each PC_macro_ (n=5).

Supplemental Table S2: Ranks the Spring OSR genotypes according to its position within each PC_macro_ (n=5).

Supplemental Table S3: Ranks the Winter OSR genotypes according to its position within each PC_alltraits_ (n=3).

Supplemental Table S4: Ranks the Spring OSR genotypes according to its position within each PC_alltraits_ (n=3).

Supplemental Table S5: Percentage of seed yield variation explained by different Partial Least Square (PLS) components for macrotraits for WOSR and SOSR.

Supplemental Table S6: Percentage of seed yield variation explained by different Partial Least Square (PLS) components for alltraits for WOSR and SOSR.

Supplemental Table S7: Partial Least Squares (PLS) regression coefficients for Winter OSR and Spring OSR for macrotraits and alltraits

Supplemental Table S8: List of 96 genotypes included in the diversity set population. The ASSYST code, genotype names, crop type description and the 4 oilseed rape groups are presented.

Supplemental Table S9: List of macrotrait (n=5) and microtrait (n=3) names and abbreviations measured in the diversity set population.

Supplemental Table S10: List of transformations applied in order to satisfy homogeneity of variances

## Acknowledgments

We thank Hannah Walpole (Rothamsted Research, UK) for help in collecting and imaging bud and ovule data and sample processing. We thank Dr. Javier Alberto Miret Barrio for his help in collecting data, harvesting and threshing the plants. Finally, we thank Amy Dodd (Rothamsted Research, UK) for the graphical representation of the oilseed rape ideotypes.

## Notes

### Competing Interest Statement

The authors have declared no competing interest.

## Literature Cited

Ahmadzadeh M, Samizadeh HA, Ahmadi MR, Soleymani F, Arantes dLC (2019) Selection Criteria for Yield Improvement in Rapeseed (Brassica napus L.). World Res. J. Agric. Sci. 6: 17–184

Angadi S, Cutforth HW, Mcconkey BG, Miller PR (2000) Response of three Brassica species to high temperature stress during reproductive growth 80:693–701

Arathi H, Ganeshaiah K, Hedge S, Ru S (1999) Seed abortion in Pongamia Pinnata (Fabaceae). American Journal of Botany 86: 659–662

Badaran R, Heravan M, Darvish F, Mahdi A (2007) Study of correlation relationships and path coefficient analysis between yield and yield components in rapeseed (Brassica napus L.). Journal of Agricultural Sciences 12: 811–819

Başalma D (2008) The correlation and path analysis of yield and yield components of different winter rapeseed (Brassica napus ssp. oleifera L.) cultivars. Research Journal of Agriculture and Biological Sciences 4: 120–125

Bennett E, Roberts JA, Wagstaff C (2012) Manipulating resource allocation in plants. J Exp Bot 63: 3391–3400

Brown JKM, Beeby R, Penfield S (2019) Yield instability of winter oilseed rape modulated by early winter temperature. Sci Rep 9: 6953

Champolivier L, Merrien A (1996) Effects of water stress applied at different growth stages to Brassica napus L. var. oleifera on yield, yield components and seed quality European Journal of Agronomy 5: 153–160.

Chen F, Zhang J, Qi C, Pu H, Chen S (2013) The analysis on diversity of germplasm resource in Brassica napus L. Jiangsu Agricultural Sciences 40: 98–99

Chen B, Xu K, Li J, Li F, Qiao J, Li H, Gao G, Yan G, Wu X (2014) Evaluation of yield and agronomic traits and their genetic variation in 488 global collections of Brassica napus L. Genetic Resources and Crop Evolution 61: 979–999

Diepenbrock W (2000) Yield analysis of winter oilseed rape (Brassica napus L.): a review. Field Crops Research 67: 35–49

Dong H, Tan C, Li Y, He Y, Wei S, Cui Y, Chen Y, Wei D, Fu Y, He Y, Wan H, Liu Z, Xiong Q, Lu K, Li J, Qian W (2018) Genome-Wide Association Study Reveals Both Overlapping and Independent Genetic Loci to Control Seed Weight and Silique Length in Brassica napus. Front Plant Sci 9: 921

Elferjani R, Soolanayakanahally R (2018) Canola Responses to Drought, Heat, and Combined Stress: Shared and Specific Effects on Carbon Assimilation, Seed Yield, and Oil Composition. Front Plant Sci 9

Food and Agriculture Organization of the United Nations (2019). FAOSTAT 2017. Available online at: http://faostat.fao.org

Habekotté B (1997) Options for increasing seed yield of winter oilseed rape (Brassica napus L.) : a simulation study. Field Crops Research 54: 109–126

Harper AL, Trick M, Higgins J, Fraser F, Clissold L, Wells R, Hattori C, Werner P, Bancroft I (2012) Associative transcriptomics of traits in the polyploid crop species Brassica napus. Nat Biotechnol 30: 798–802

Havlickova L, He Z, Wang L, Langer S, Harper AL, Kaur H, Broadley MR, Gegas V, Bancroft I (2018) Validation of an updated Associative Transcriptomics platform for the polyploid crop species Brassica napus by dissection of the genetic architecture of erucic acid and tocopherol isoform variation in seeds. Plant J 93: 181–192

Hu Q, Hua W, Yin Y, Zhang X, Liu L, Shi J, Zhao Y, Qin L, Chen C, Wang H (2017) Rapeseed research and production in China. The Crop Journal 5: 127–135

Kenward MG, Roger JH (1997) Small sample inference for fixed effects from restricted maximum likelihood. Biometrics 53: 983–997.

Kuai J, Sun Y, Zuo Q, Huang H, Liao Q, Wu C, Lu J, Wu J, Zhou G (2015) The yield of mechanically harvested rapeseed (Brassica napus L.) can be increased by optimum plant density and row spacing. Sci Rep 5: 18835

Lardon A, Triboi-Blondel A-M (1995) Cold and freezer stress at flowering-effects on seed yield in winter rapeseed. Field Crops Res 44, 95–101.

Li N, Peng W, Shi J, Wang X, Liu G, Wang H (2015) The Natural Variation of Seed Weight Is Mainly Controlled by Maternal Genotype in Rapeseed (Brassica napus L.). PLoS One 10: e0125360

Li N, Song D, Peng W, Zhan J, Shi J, Wang X, Liu G, Wang H (2019) Maternal control of seed weight in rapeseed (Brassica napus L.): the causal link between the size of pod (mother, source) and seed (offspring, sink). Plant Biotechnol J 17: 736–749

Moradi M, Hoveize M, Shahbazi E (2017) Study the relations between grain yield and related traits in canola y multivariate analysis. Journal of Crop Breeding 9: 187–194

Naazar A, Javidfar F, Elmira JY, Mirza MY (2003) Relationship among yield components and selection criteria for yield improvement in winter rapeseed (Brassica napus L.). Pak. J. Bot., 35: 167–174

Nesi N, Delourme R, Bregeon M, Falentin C, Renard M (2008) Genetic and molecular approaches to improve nutritional value of Brassica napus L. seed. C R Biol 331: 763–771

Özer H, Oral E, Dogru U (1999) Relationships between yield and yield components on currently improved spring rapeseed cultivars. Tr. J. of Agriculture and Forestry: 603–607

Pinet A, Mathieu A, Jullien A (2015) Floral bud damage compensation by branching and biomass allocation in genotypes of Brassica napus with different architecture and branching potential. Front Plant Sci 6: 70

Raboanatahiry N, Chao H, Dalin H, Pu S, Yan W, Yu L, Wang B, Li M (2018) QTL Alignment for Seed Yield and Yield Related Traits in Brassica napus. Front Plant Sci 9: 1127

Ren T, Liu B, Lu J, Deng Z, Li X, Cong R (2017) Optimal plant density and N fertilization to achieve higher seed yield and lower N surplus for winter oilseed rape (Brassica napus L.). Field Crops Research 204: 199–207

Rohart F, Gautier B, Singh A, Le Cao KA (2017) mixOmics: An R package for ‘omics feature selection and multiple data integration. PLoS Comput Biol 13: e1005752

Sabaghnia N, Dehghani H, Mabm (2010) Interrelationships between seed yield and 20 related traits of 49 canola (Brassica napus L.) genotypes in non-stressed and water-stressed environments. Spanish Journal of Agricultural Research 8: 356–370

Sadras VO (2007) Evolutionary aspects of the trade-off between seed size and number in crops. Field Crops Research 100: 125–138

Schiessl S, Iniguez-Luy F, Qian W, Snowdon RJ (2015) Diverse regulatory factors associate with flowering time and yield responses in winter-type Brassica napus. BMC Genomics 16: 737

Shi J, Li R, Qiu D, Jiang C, Long Y, Morgan C, Bancroft I, Zhao J, Meng J (2009) Unraveling the complex trait of crop yield with quantitative trait loci mapping in Brassica napus. Genetics 182: 851–861

Siles L, Eastmond P, Kurup S (2020) Big data from small tissues: extraction of high-quality RNA for RNA-sequencing from different oilseed Brassica seed tissues during seed development. Plant Methods 16: 80

Snowdon R, Wilfried L, Friedt W (2007) Genome Mapping and Molecular Breeding in Plants, Volume 2 Oilseeds. In: C. Kole (Ed.)

Stahl A, Vollrath P, Samans B, Frisch M, Wittkop B, Snowdon RJ (2019) Effect of breeding on nitrogen use efficiency-associated traits in oilseed rape. Journal of Experimental Botany 70: 1969–1986

Tariq H, Tanveer SK, Qamar M, Javaid RA, Vaseer SG, Jhanzab HM, Hassan MJ, Iqbal H (2020) Correlation and path analysis of Brassica napus genotypes for yield related traits. Life Science Journal 17

Tunçtürk M, Çiçti V (2007) Relationships between yield and some yield components in rapeseed (Brassica napus ssp. Oleifera L.) cultivars by using correlation and path analysis. Pak. J. Bot. 39: 81–84

Ul-Hasan E, Mustafa HSB, Bibi T, Mahmood T (2014) Genetic Variability, correlation and path analysis in advanced lines of rapeseed (Brassica napus L.) for yield components. Cercetari Agronomice in Moldova 47: 71–79

Wang X, Mathieu A, Cournède P-H, Allirand J-M, Jullien A, de Reffye P, Zhang BG (2011) Variability and regulation of the number of ovules, seeds and pods according to assimilate availability in winter oilseed rape (Brassica napus L.). Field Crops Research 122: 60–69

Welham SJ, Gezan S A, Clark S J, Mead A (2015) Statistical methods in biology: design and analysis of experiments and regression. CRC Press, Boca Raton, Florida.

Weymann W, Böttcher U, Sieling K, Kage H (2015) Effects of weather conditions during different growth phases on yield formation of winter oilseed rape. Field Crops Research 173: 41–48

Yang Y, Shi J, Wang X, Liu G, Wang H (2016) Genetic architecture and mechanism of seed number per pod in rapeseed: elucidated through linkage and near-isogenic line analysis. Sci Rep 6: 24124

Yang Y, Wang Y, Zhan J, Shi J, Wang X, Liu G, Wang H (2017) Genetic and cytological analyses of the natural variation of seed number per pod in rapeseed (Brassica napus L.). Front Plant Sci 8: 1890

Yu K, Wang X, Chen F, Chen S, Peng Q, Li H, Zhang W, Hu M, Chu P, Zhang J, Guan R (2016) Genome-wide transcriptomic analysis uncovers the molecular basis underlying early flowering and apetalous characteristic in Brassica napus L. Sci Rep 6: 30576.

Young LW, Wilen RW, Bonham-Smith PC. High temperature stress of Brassica napus during flowering reduces micro- and megagametophyte fertility, induces fruit abortion, and disrupts seed production. Journal of Experimental Botany 55: 485–495.

Zhang L, Yang G, Liu P, Hong D, Li S, He Q (2011) Genetic and correlation analysis of silique-traits in Brassica napus L. by quantitative trait locus mapping. Theor Appl Genet 122: 21–31

Zhu Y, Ye J, Zhan J, Zheng X, Zhang J, Shi J, Wang X, Liu G, Wang H (2020) Validation and characterization of a seed number per silique quantitative trait locus qSN.A7 in Rapeseed (Brassica napus L.). Front Plant Sci 11: 68

