## Supplementary figures and images for "Uncovering the ideal plant ideotype for maximising seed yield in *Brassica napus*"

### Supplemental Figure S1

#### Slide 1
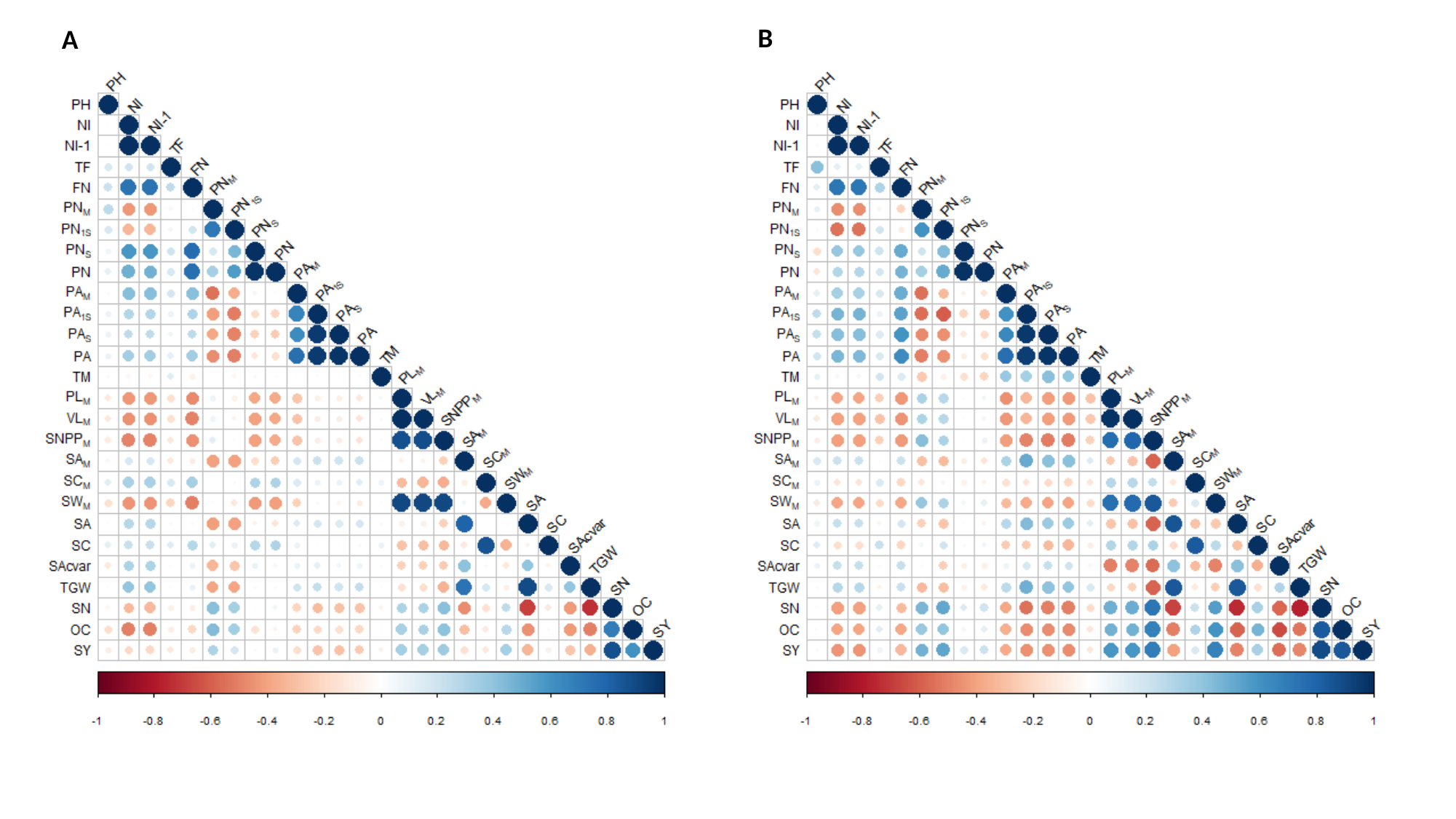

B
A

### Supplemental Figure S2

#### Slide 1
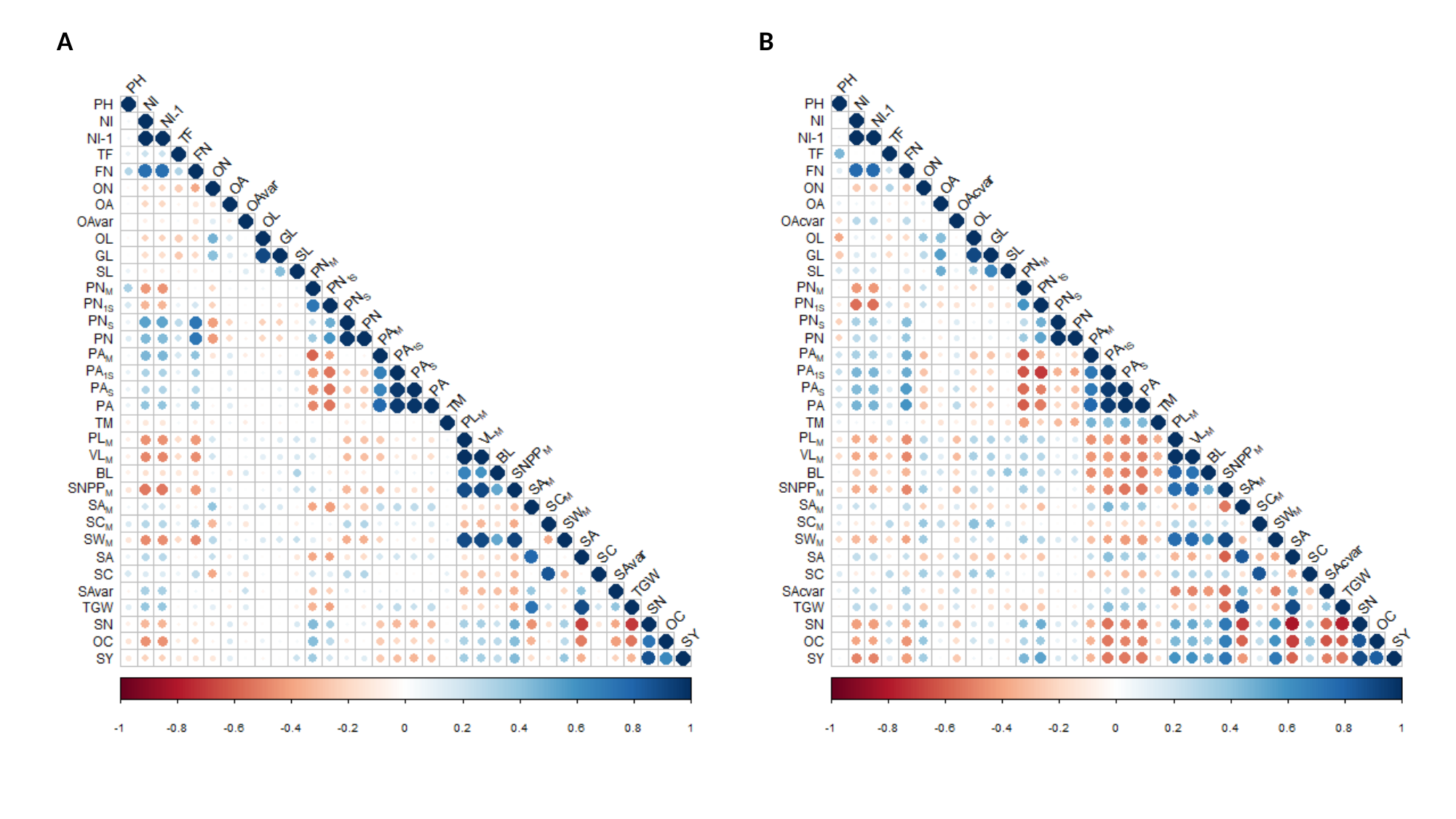

A
B

### Supplemental File S1

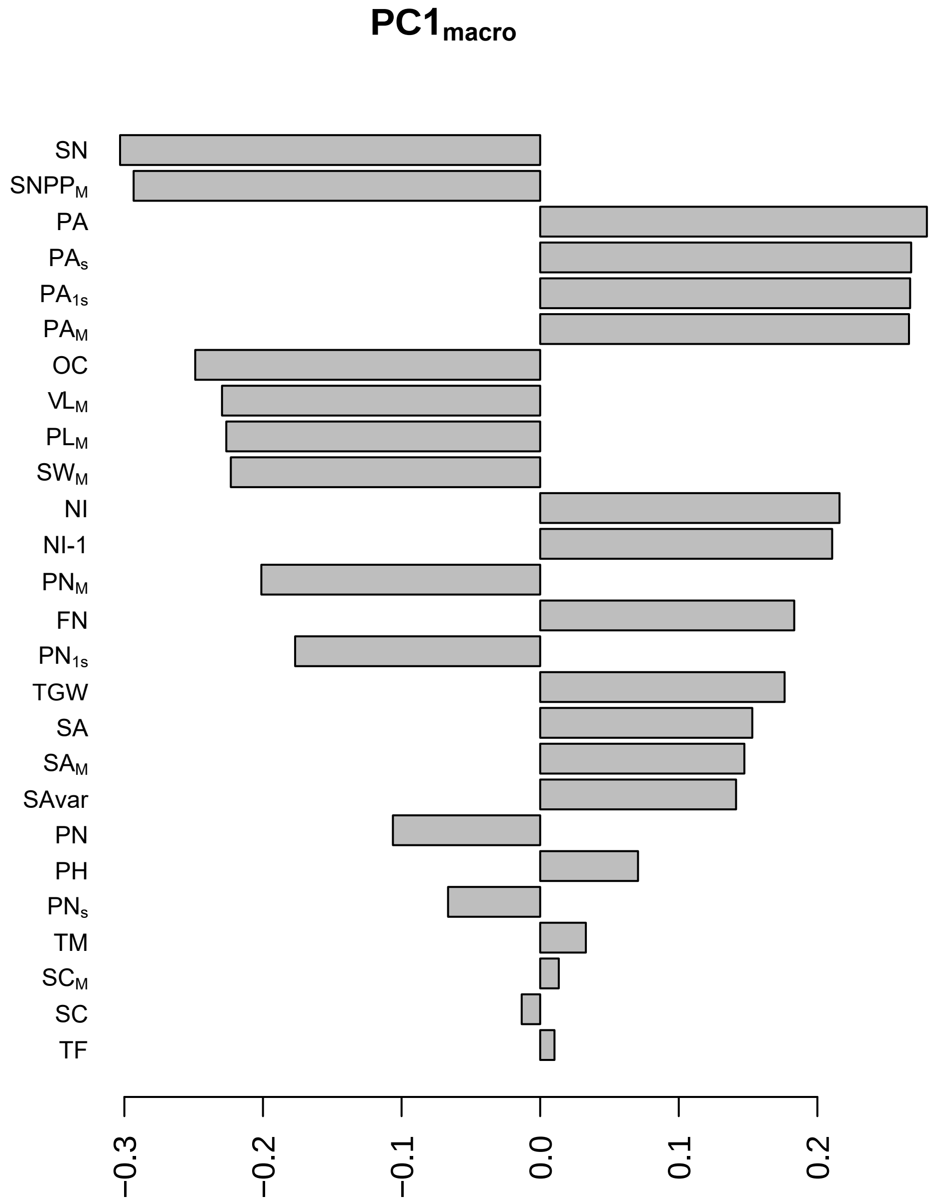

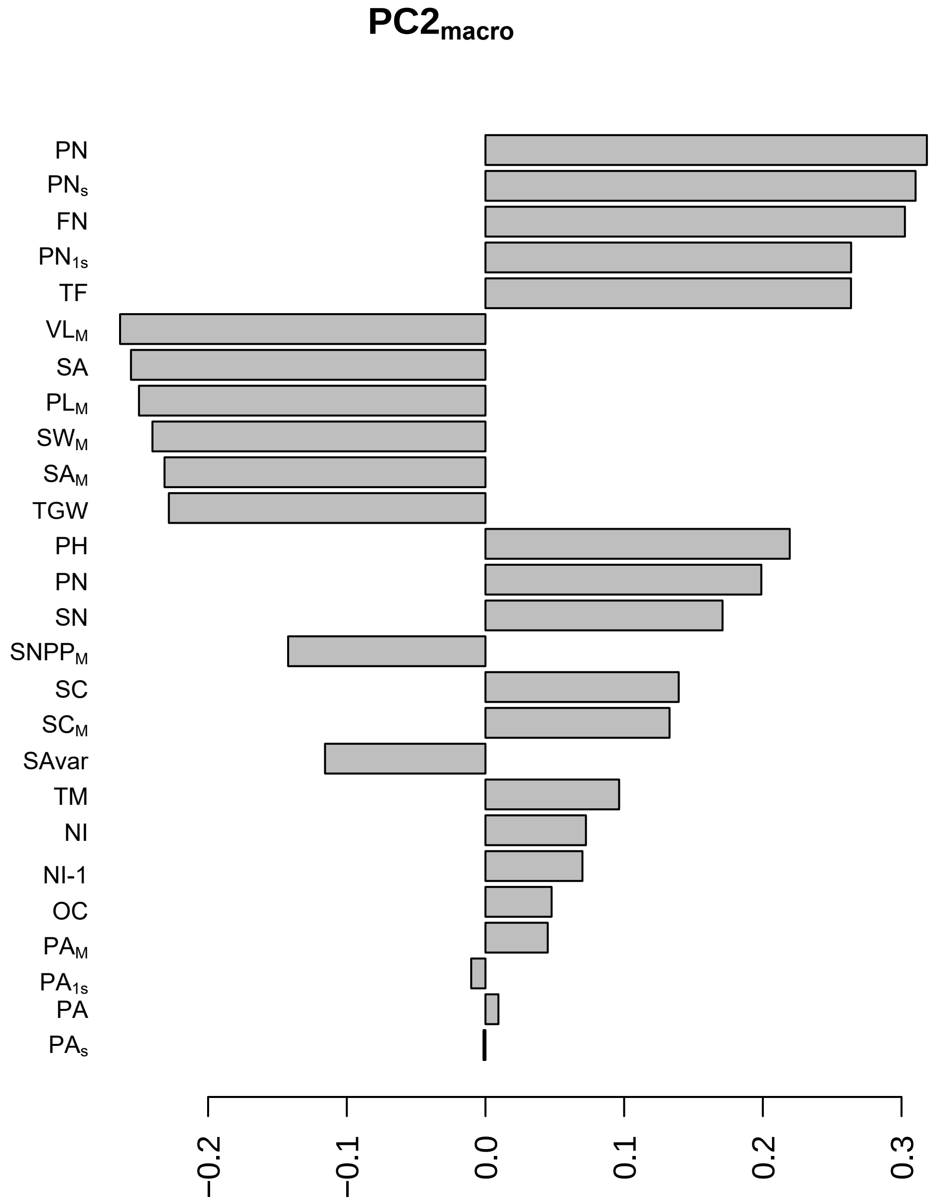

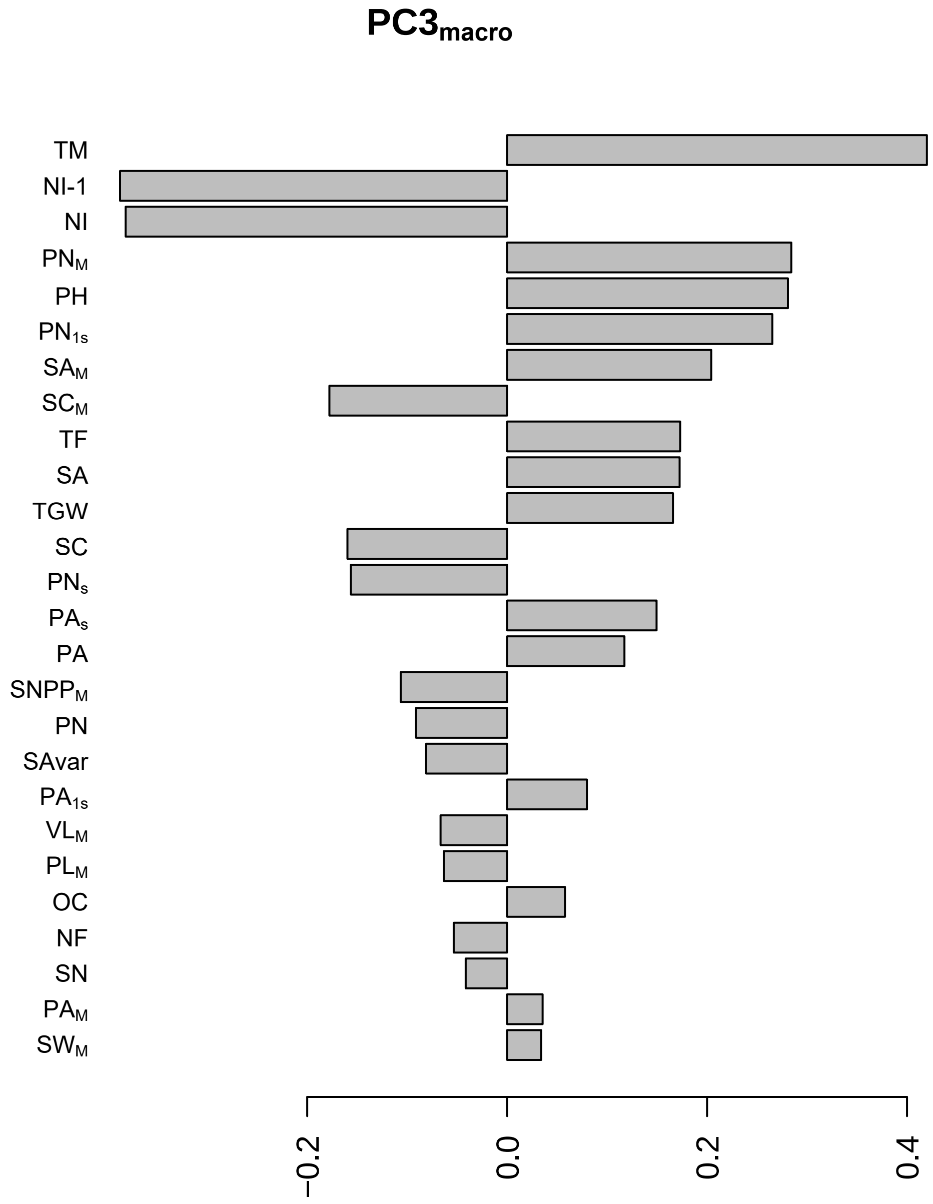

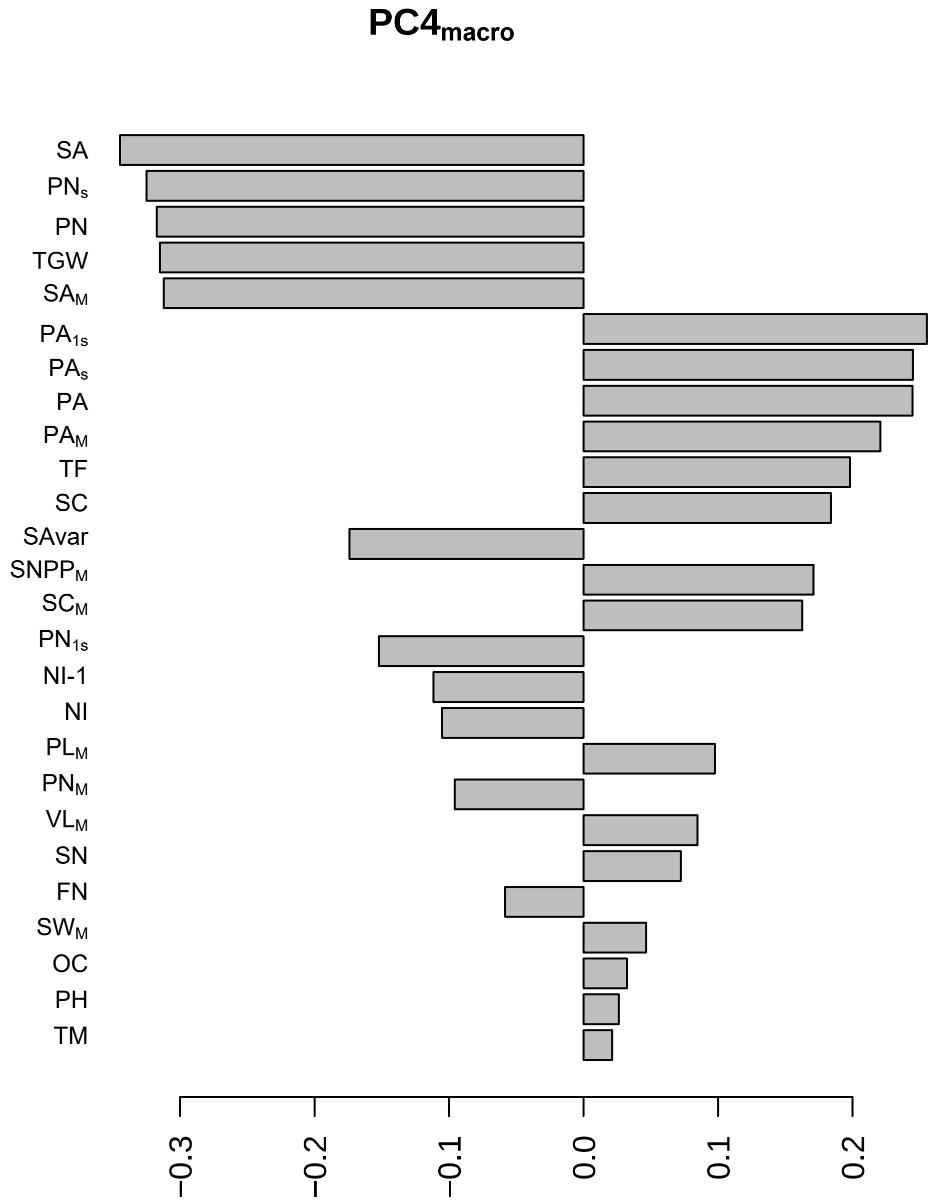

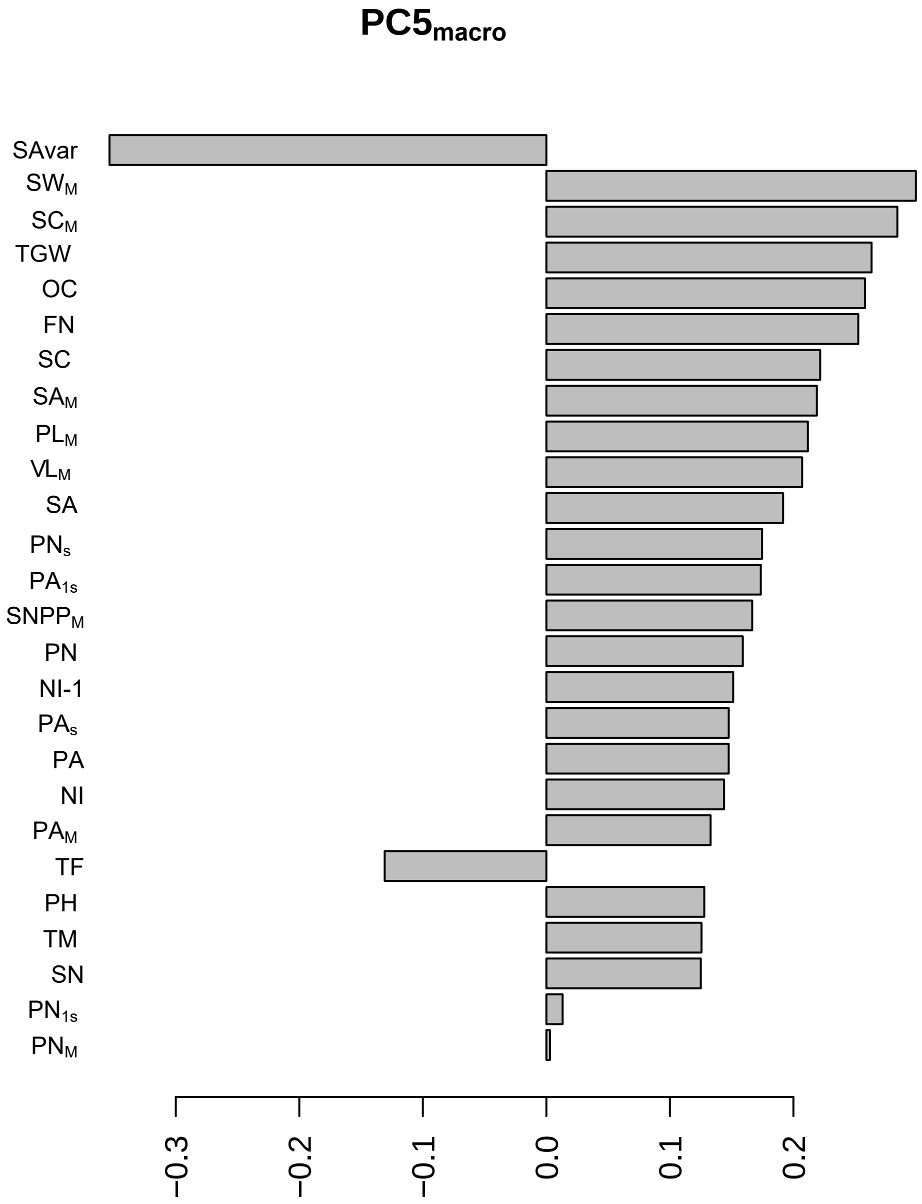

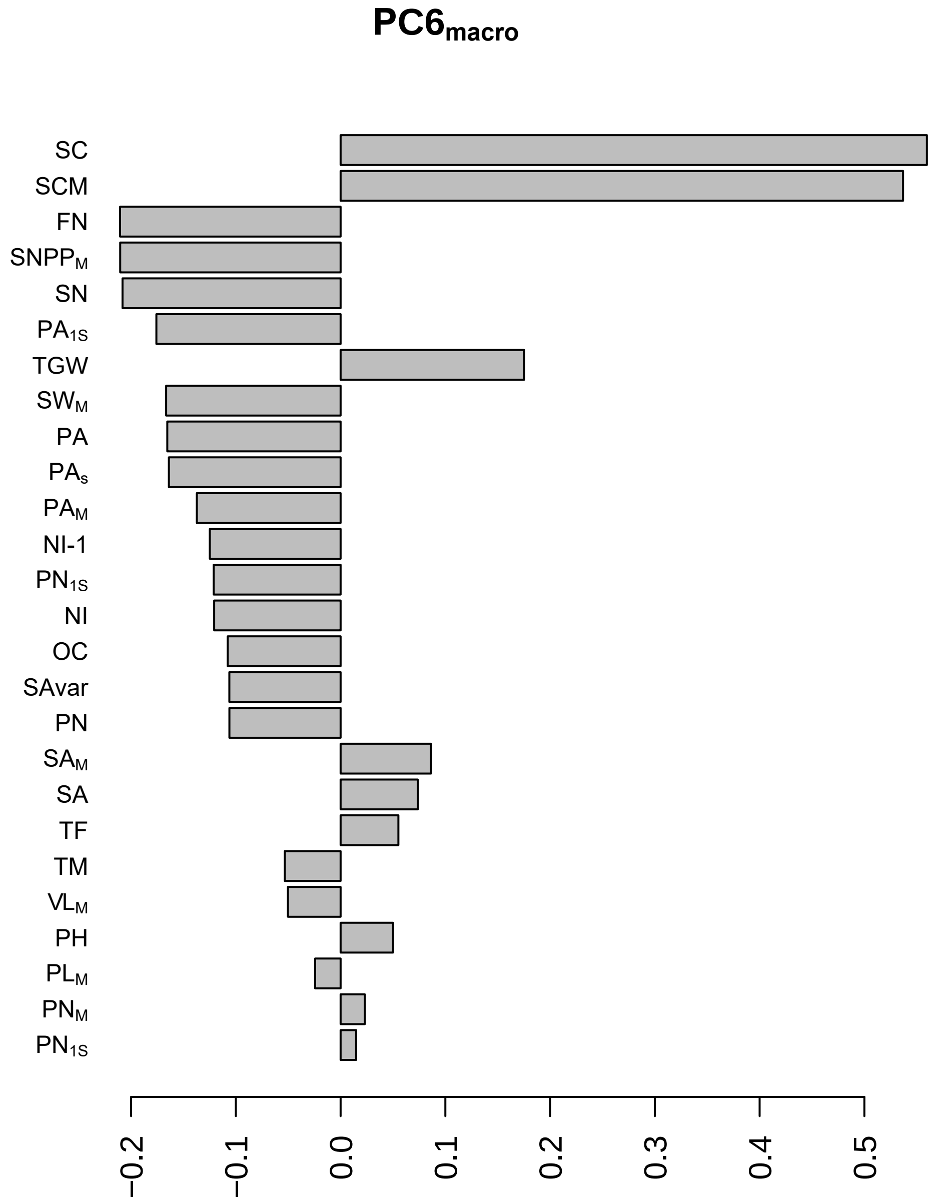

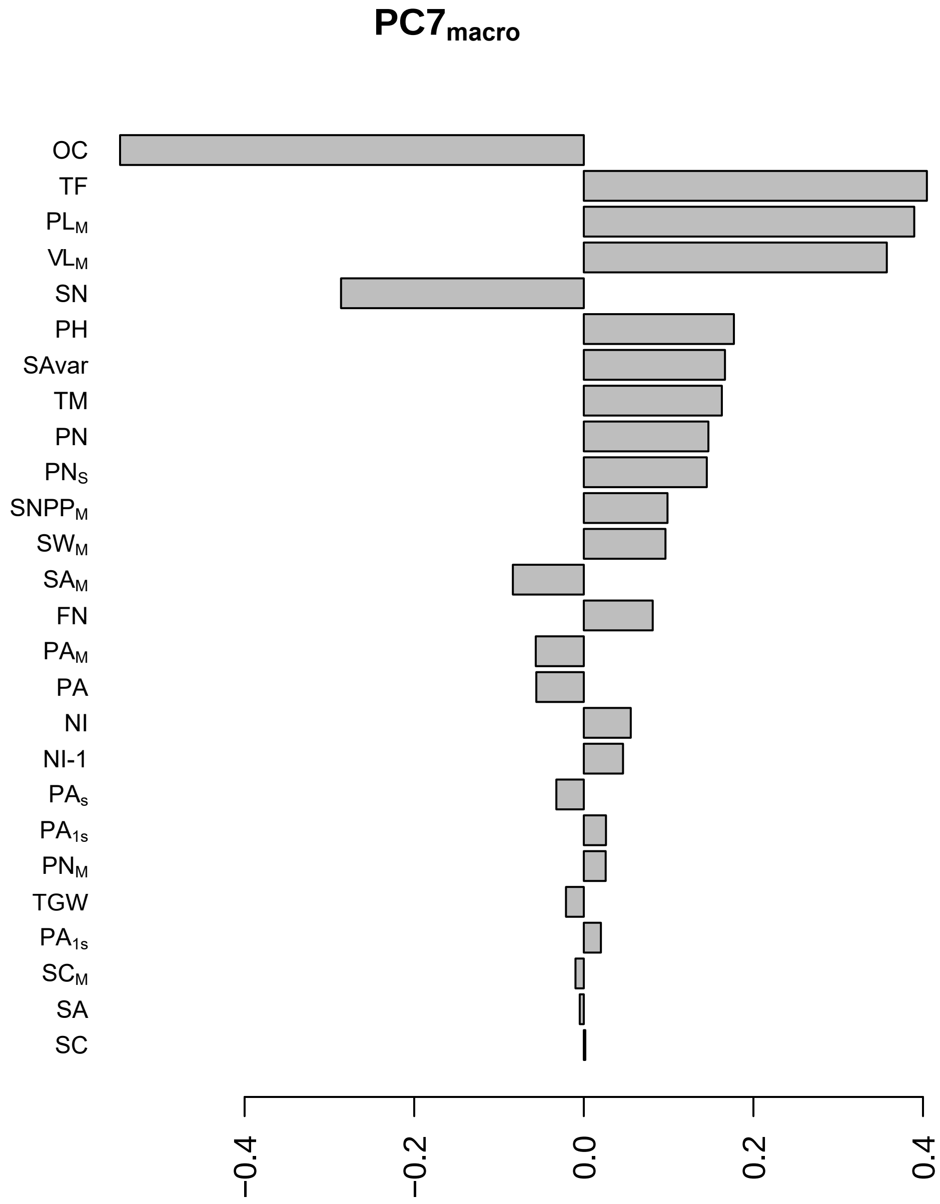

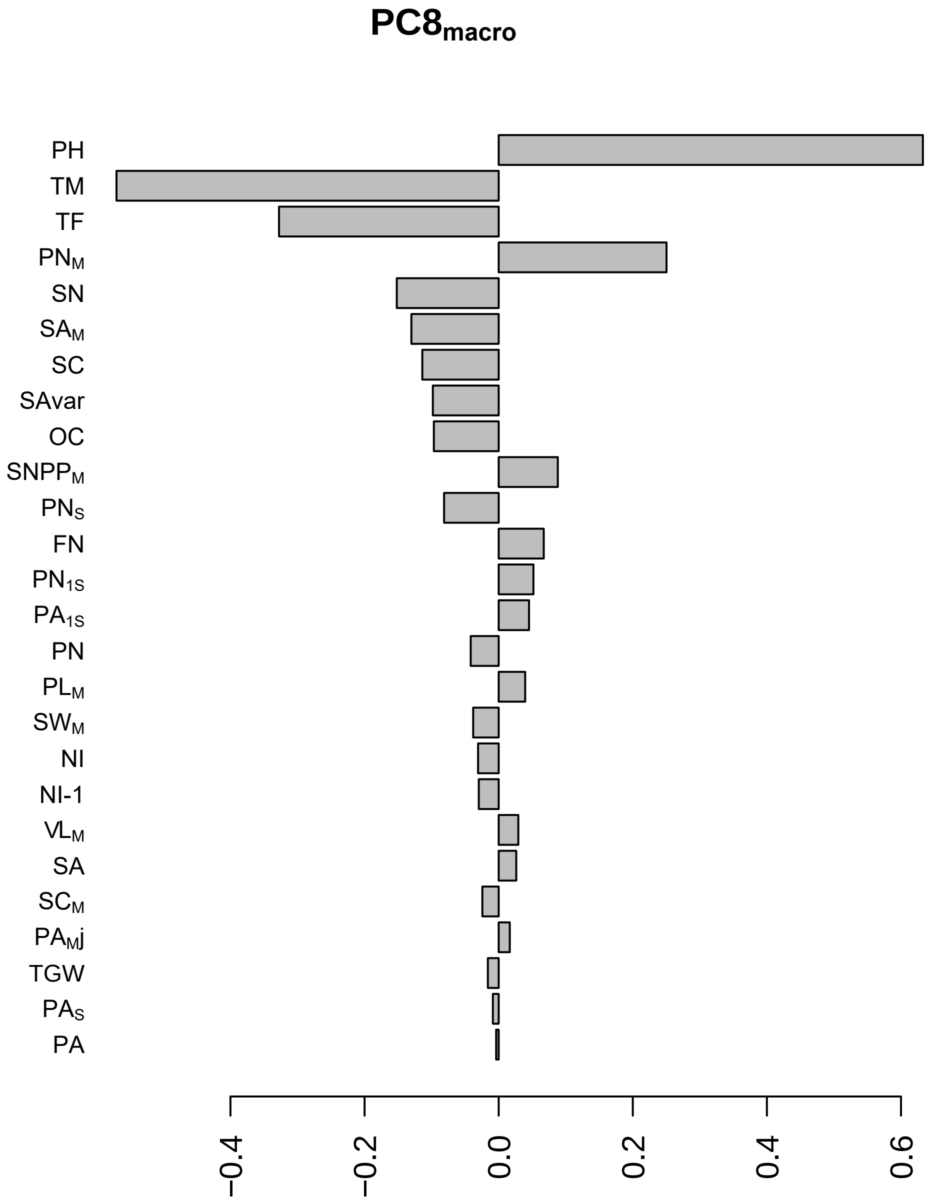

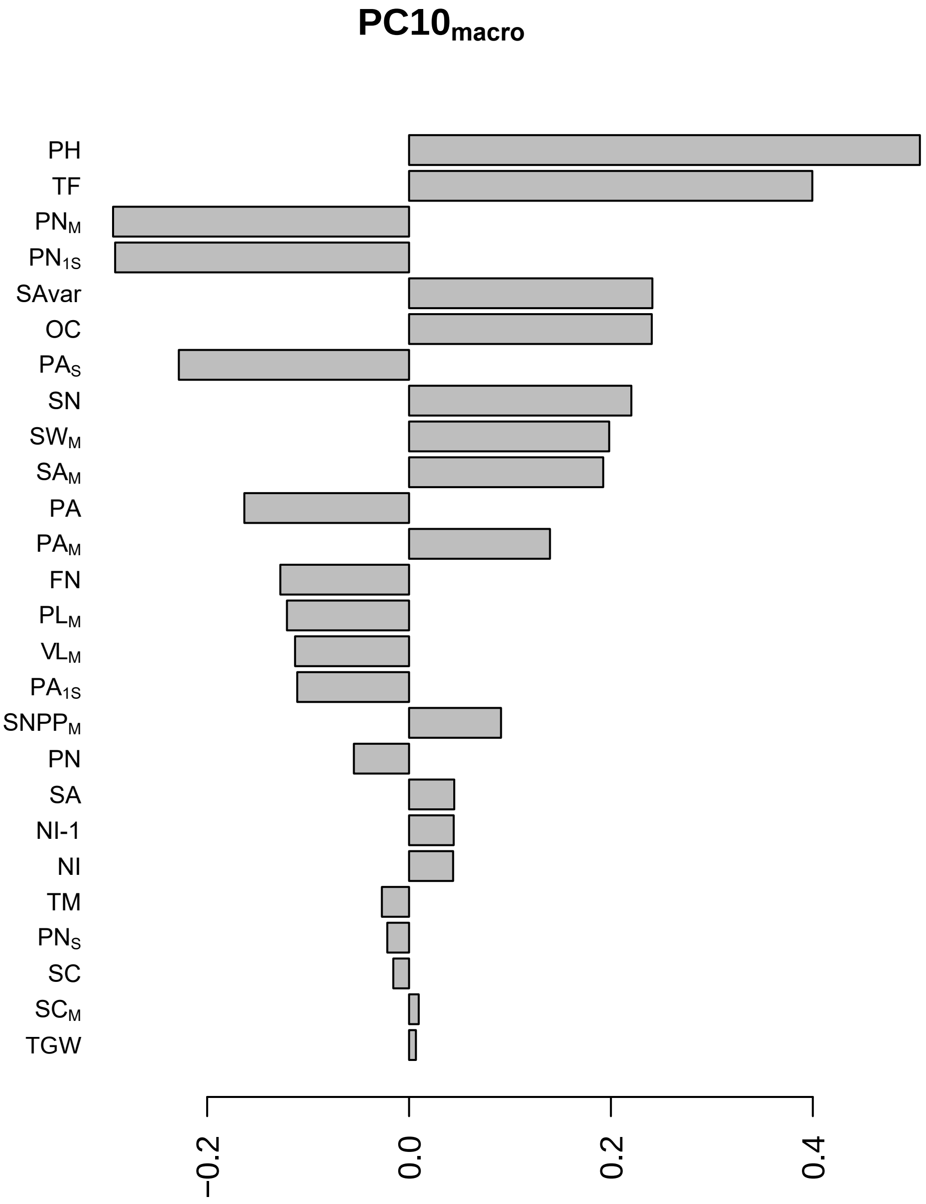

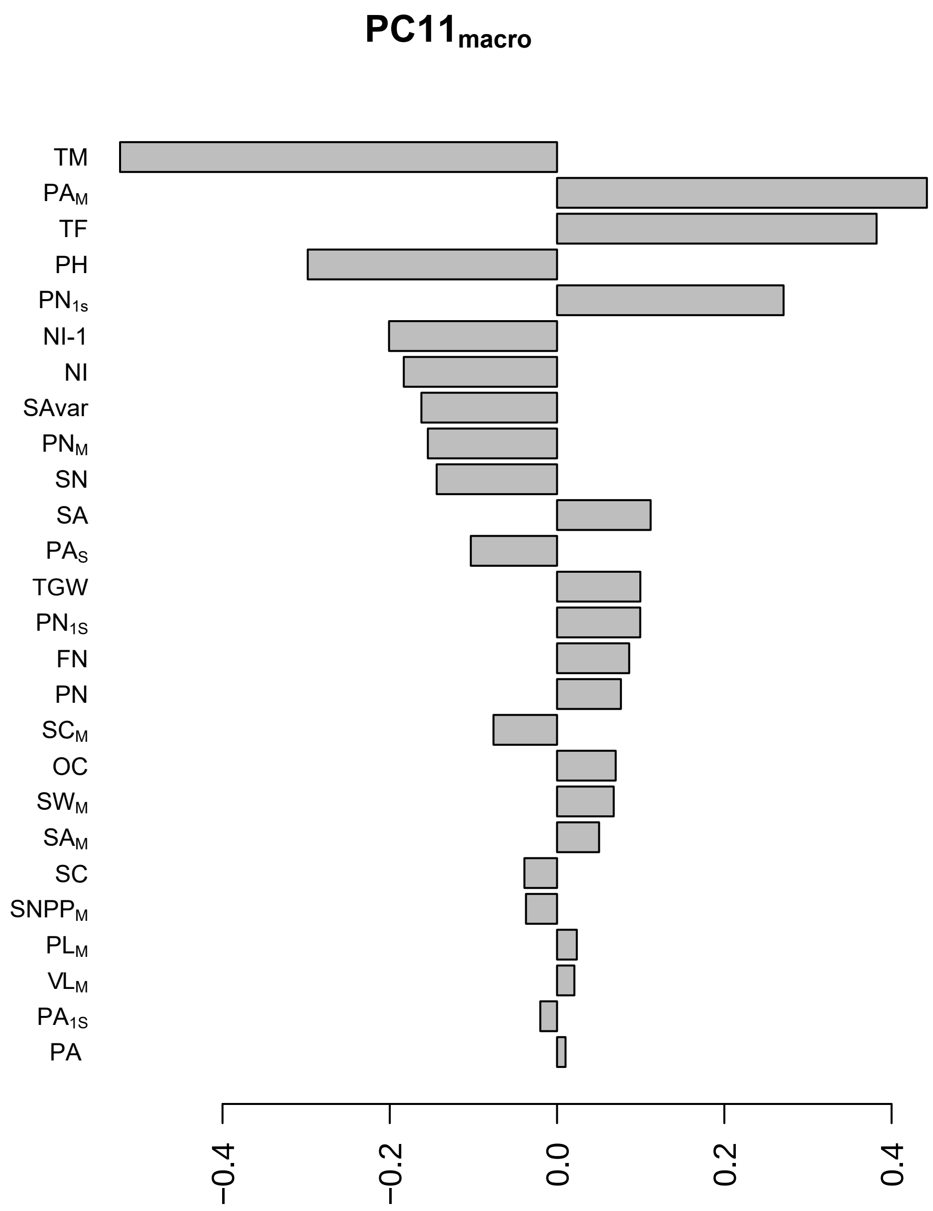
