## Supplemental File S2 for "Uncovering the ideal plant ideotype for maximising seed yield in *Brassica napus*"

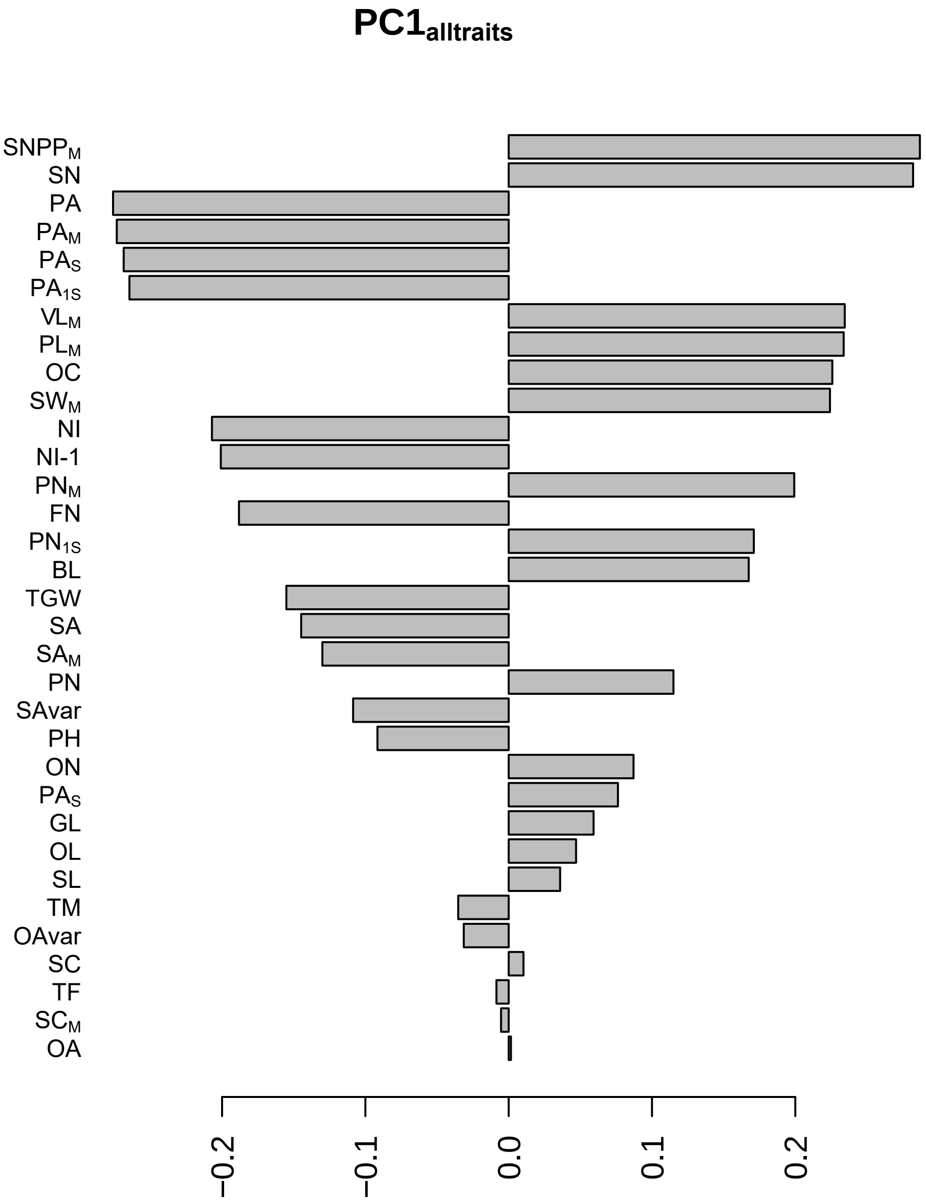

PC1alltraits
SNPPM
SN
PA
PAM
PAS
PA1S
VLM
PLM
OC
SWM
NI
NI-1
PNM
FN
PN1S
BL
TGW
SA
SAM
PN
SAvar
PH
ON
PAS
GL
OL
SL
TM
OAvar
SC
TF
SCM
OA
−0.2
−0.1
0.0
0.1
0.2

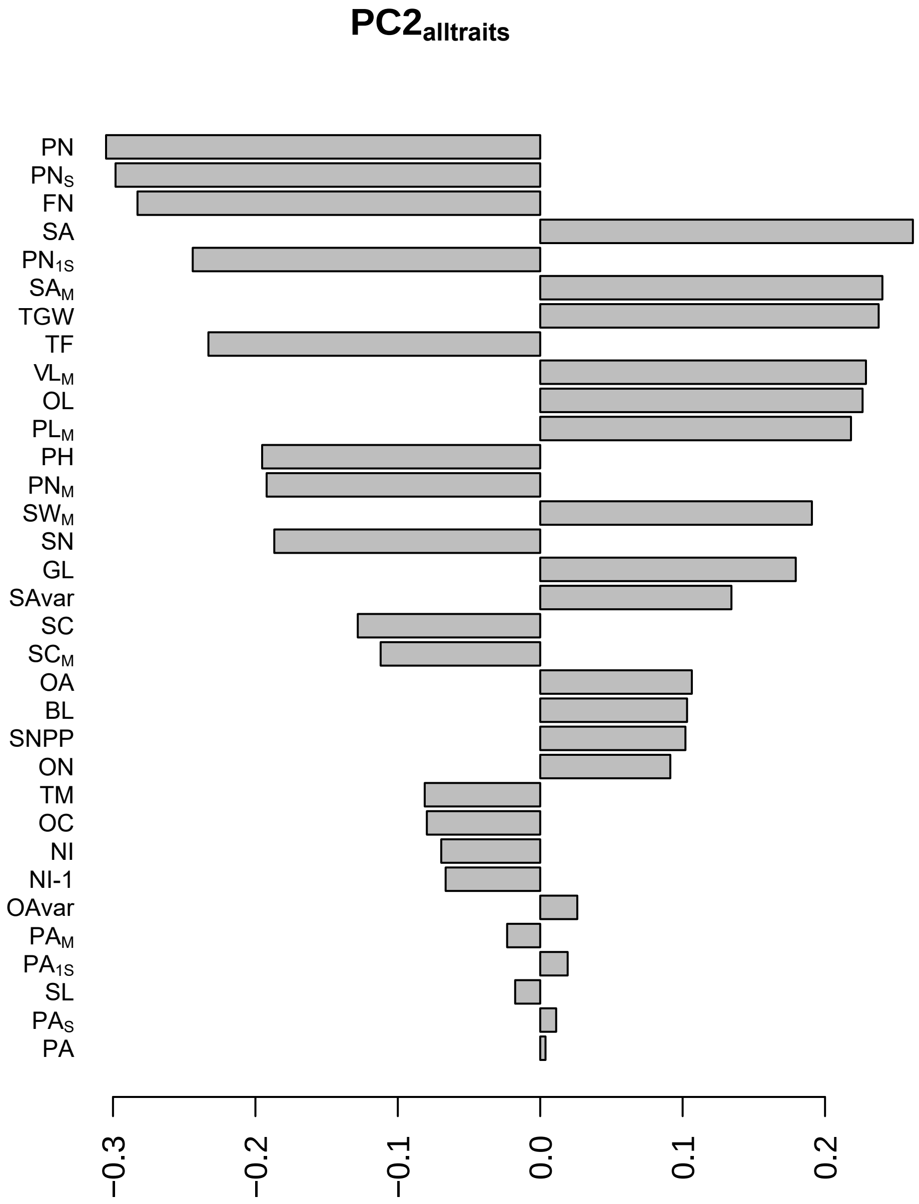
PC2alltraits
PN
PNS
FN
SA
PN1S
SAM
TGW
TF
VLM
OL
PLM
PH
PNM
SWM
SN
GL
SAvar
SC
SCM
OA
BL
SNPP
ON
TM
OC
NI
NI-1
OAvar
PAM
PA1S
SL
PAS
PA
−0.3
−0.2
−0.1
0.0
0.1
0.2

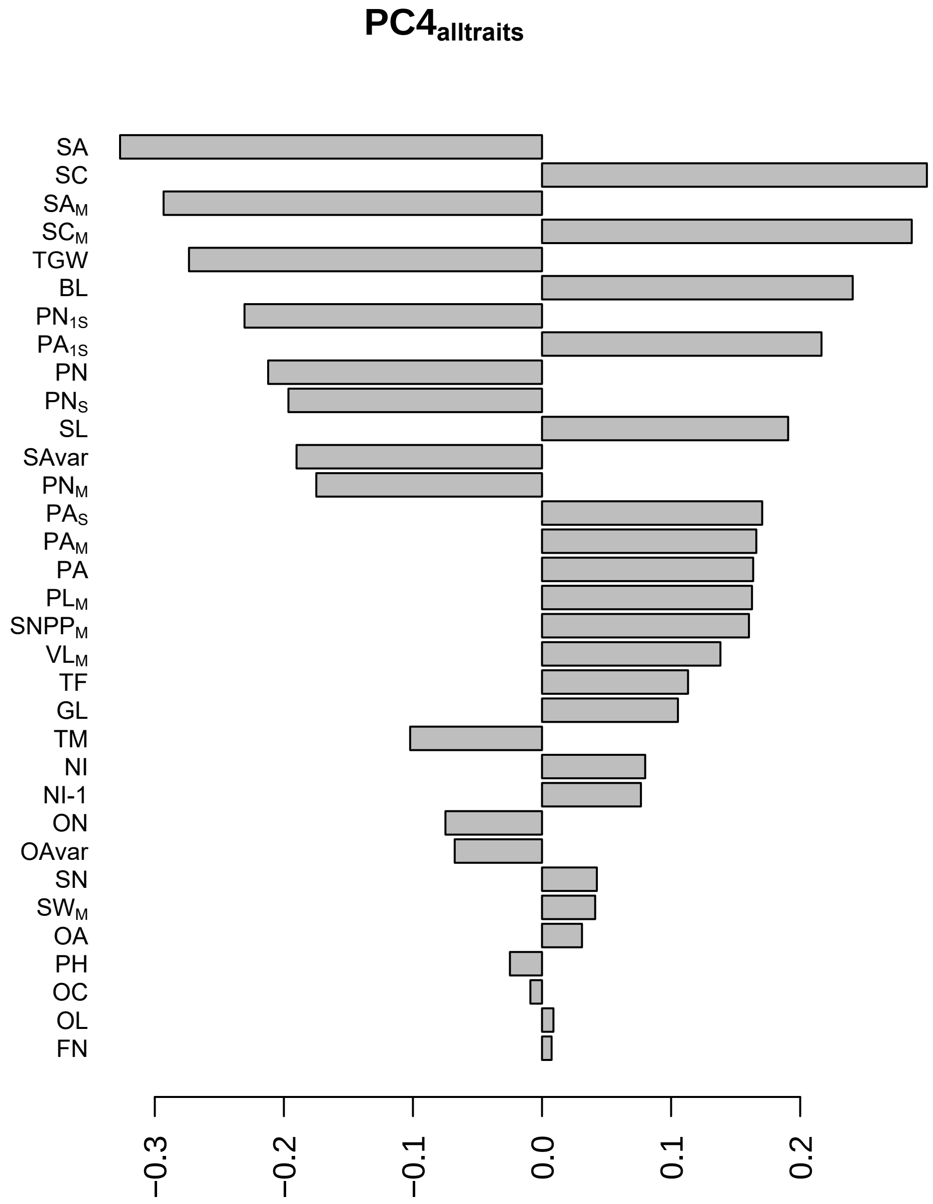

PC4alltraits
SA
SC
SAM
SCM
TGW
BL
PN1S
PA1S
PN
PNS
SL
SAvar
PNM
PAS
PAM
PA
PLM
SNPPM
VLM
TF
GL
TM
NI
NI-1
ON
OAvar
SN
SWM
OA
PH
OC
OL
FN
−0.3
−0.2
−0.1
0.0
0.1
0.2

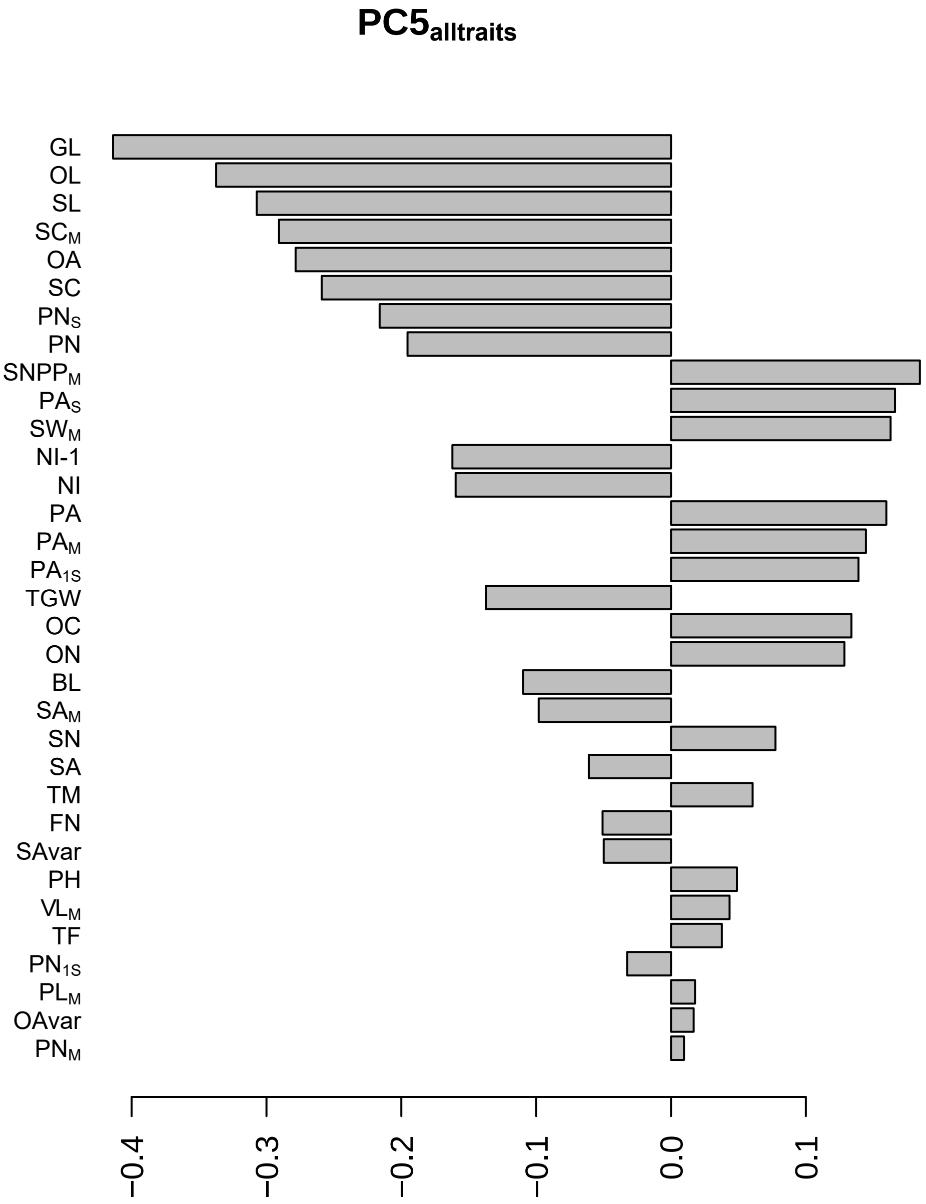
PC5alltraits
GL
OL
SL
SCM
OA
SC
PNS
PN
SNPPM
PAS
SWM
NI-1
NI
PA
PAM
PA1S
TGW
OC
ON
BL
SAM
SN
SA
TM
FN
SAvar
PH
VLM
TF
PN1S
PLM
OAvar
PNM
−0.4
−0.3
−0.2
−0.1
0.0
0.1

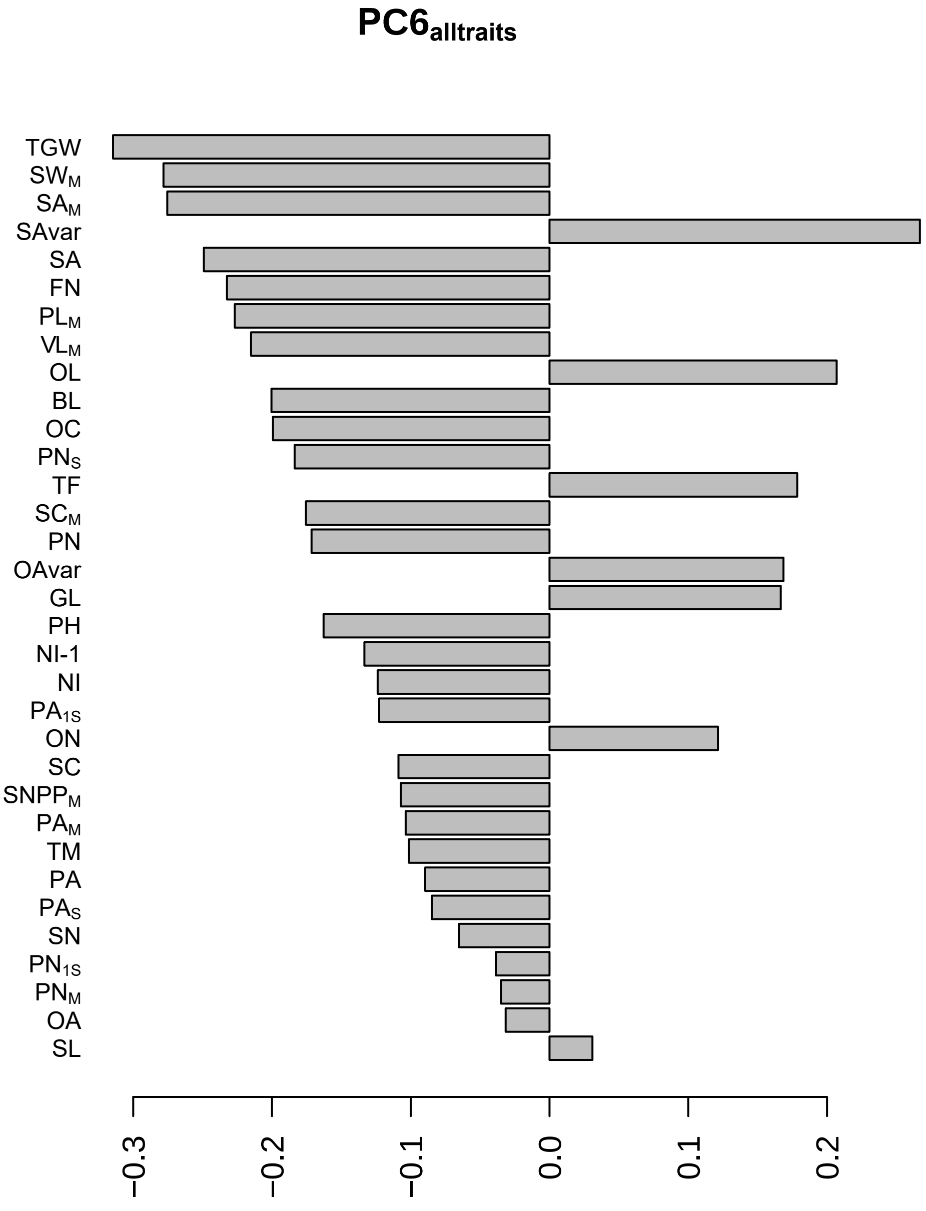

PC6alltraits
TGW
SWM
SAM
SAvar
SA
FN
PLM
VLM
OL
BL
OC
PNS
TF
SCM
PN
OAvar
GL
PH
NI-1
NI
PA1S
ON
SC
SNPPM
PAM
TM
PA
PAS
SN
PN1S
PNM
OA
SL
−0.3
−0.2
−0.1
0.0
0.1
0.2

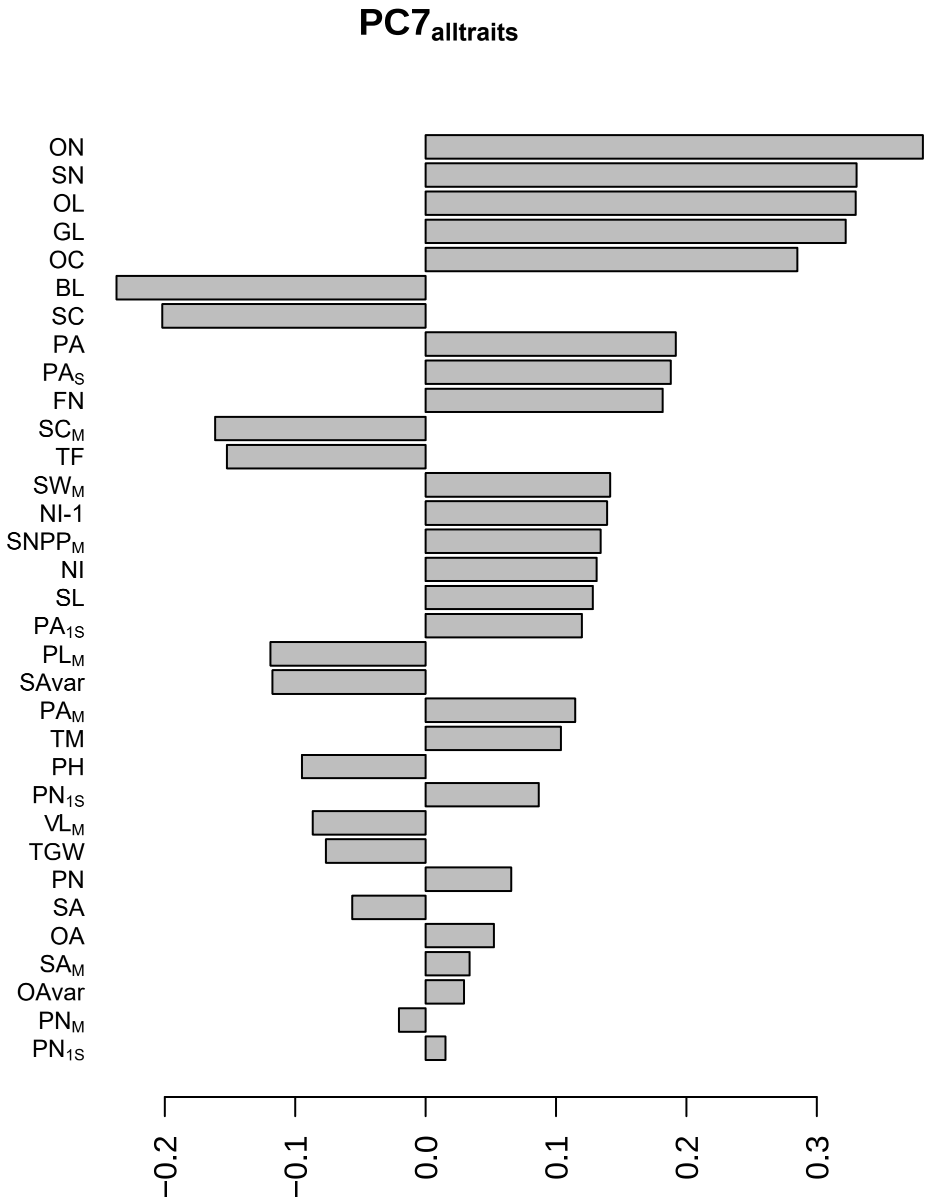

PC7alltraits
ON
SN
OL
GL
OC
BL
SC
PA
PAS
FN
SCM
TF
SWM
NI-1
SNPPM
NI
SL
PA1S
PLM
SAvar
PAM
TM
PH
PN1S
VLM
TGW
PN
SA
OA
SAM
OAvar
PNM
PN1S
−0.2
−0.1
0.0
0.1
0.2
0.3

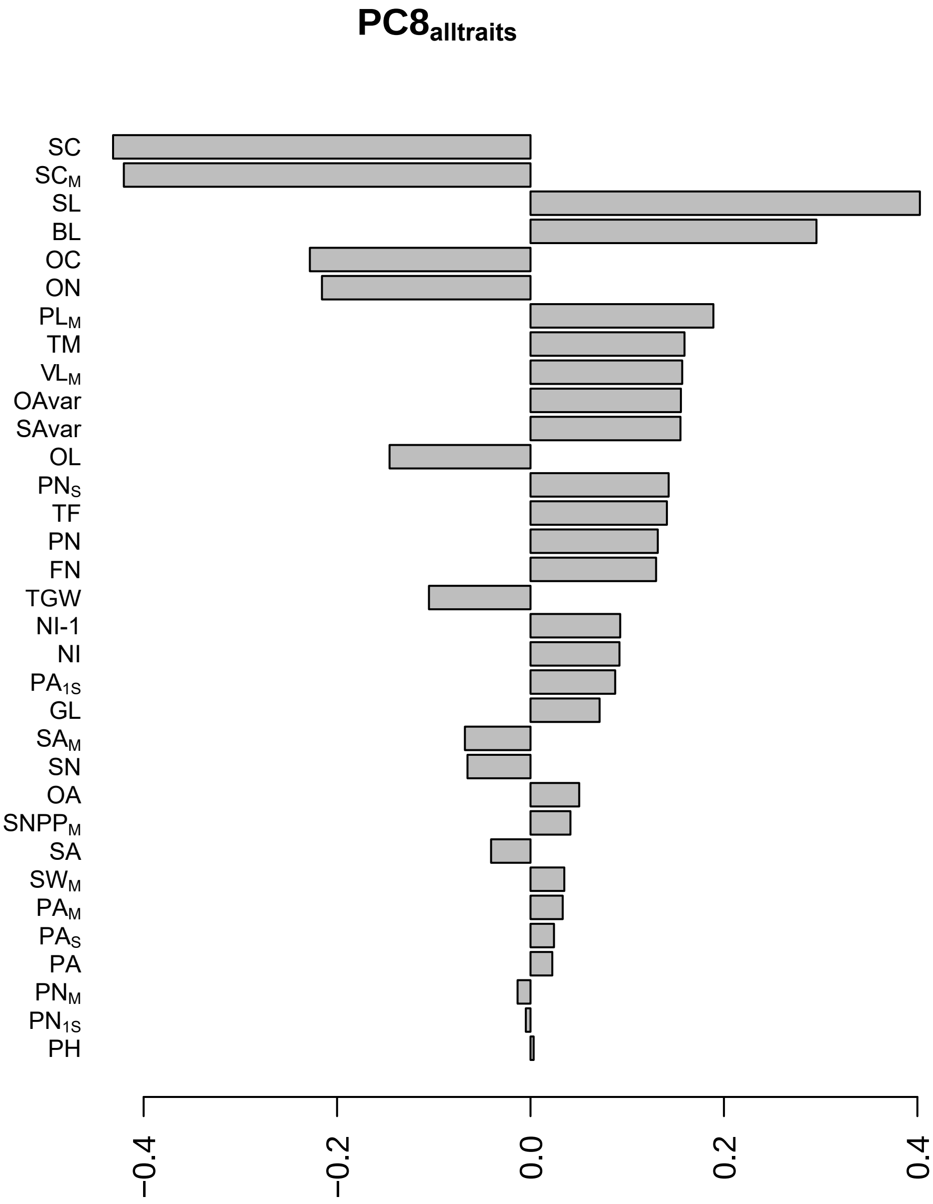

PC8alltraits
SC
SCM
SL
BL
OC
ON
PLM
TM
VLM
OAvar
SAvar
OL
PNS
TF
PN
FN
TGW
NI-1
NI
PA1S
GL
SAM
SN
OA
SNPPM
SA
SWM
PAM
PAS
PA
PNM
PN1S
PH
−0.4
−0.2
0.0
0.2
0.4

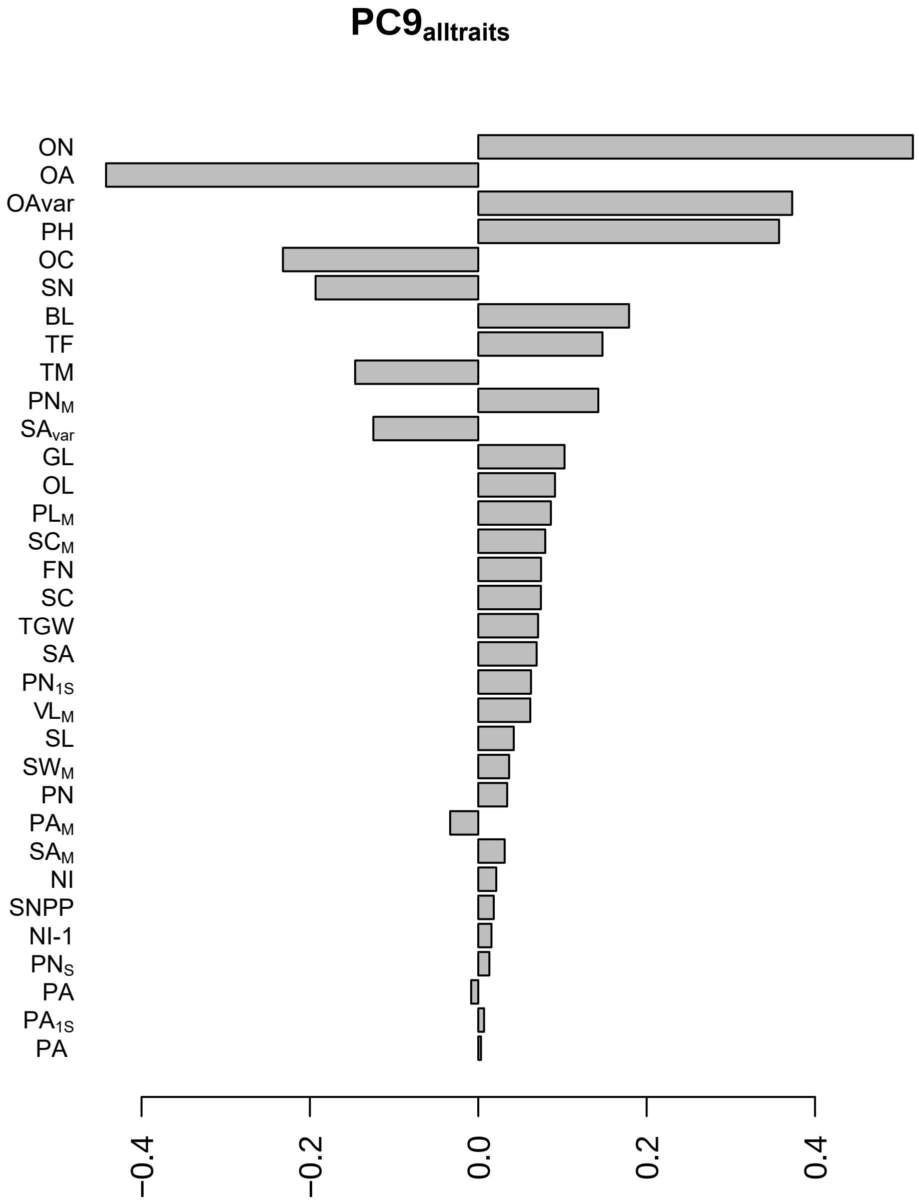

PC9alltraits
ON
OA
OAvar
PH
OC
SN
BL
TF
TM
PNM
SAvar
GL
OL
PLM
SCM
FN
SC
TGW
SA
PN1S
VLM
SL
SWM
PN
PAM
SAM
NI
SNPP
NI-1
PNS
PA
PA1S
PA
−0.4
−0.2
0.0
0.2
0.4

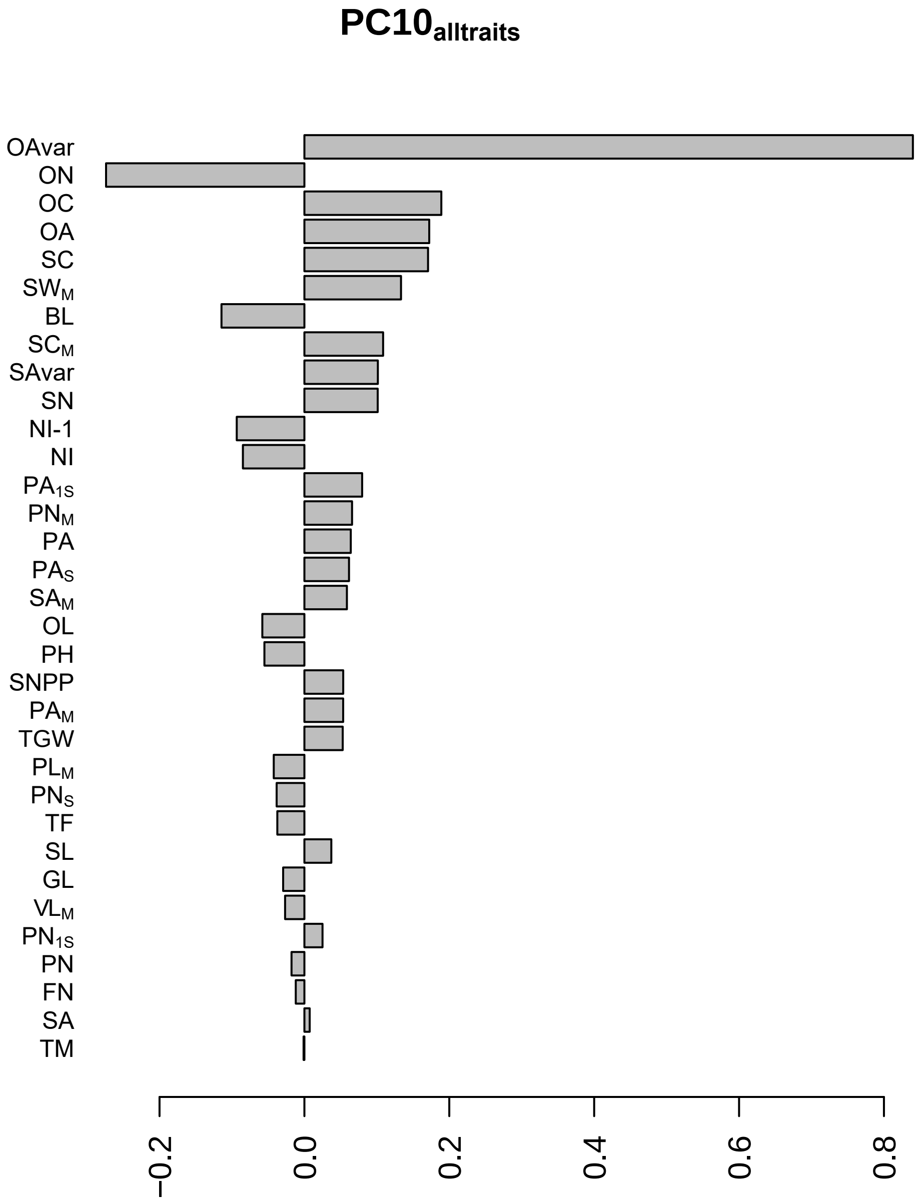
PC10alltraits
OAvar
ON
OC
OA
SC
SWM
BL
SCM
SAvar
SN
NI-1
NI
PA1S
PNM
PA
PAS
SAM
OL
PH
SNPP
PAM
TGW
PLM
PNS
TF
SL
GL
VLM
PN1S
PN
FN
SA
TM
−0.2
0.0
0.2
0.4
0.6
0.8

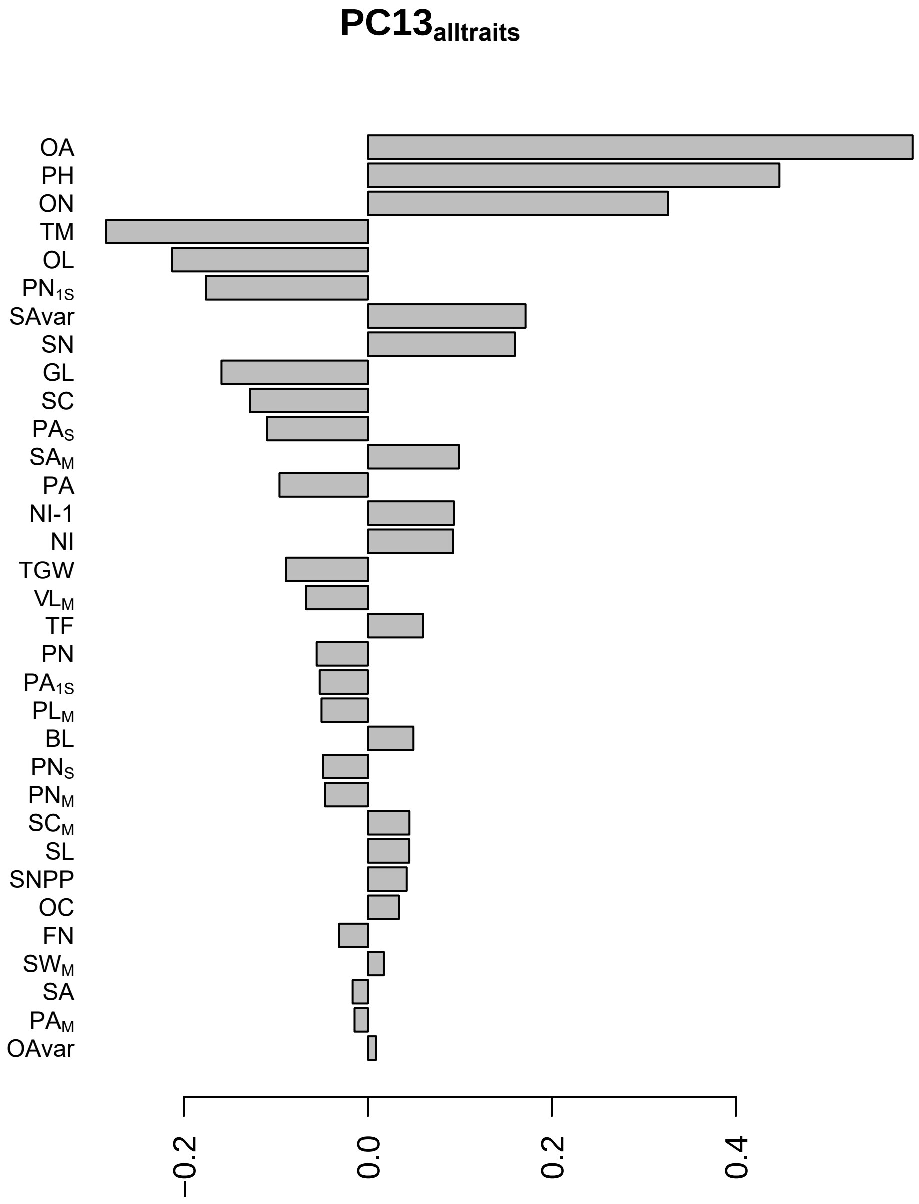

PC13alltraits
OA
PH
ON
TM
OL
PN1S
SAvar
SN
GL
SC
PAS
SAM
PA
NI-1
NI
TGW
VLM
TF
PN
PA1S
PLM
BL
PNS
PNM
SCM
SL
SNPP
OC
FN
SWM
SA
PAM
OAvar
−0.2
0.0
0.2
0.4

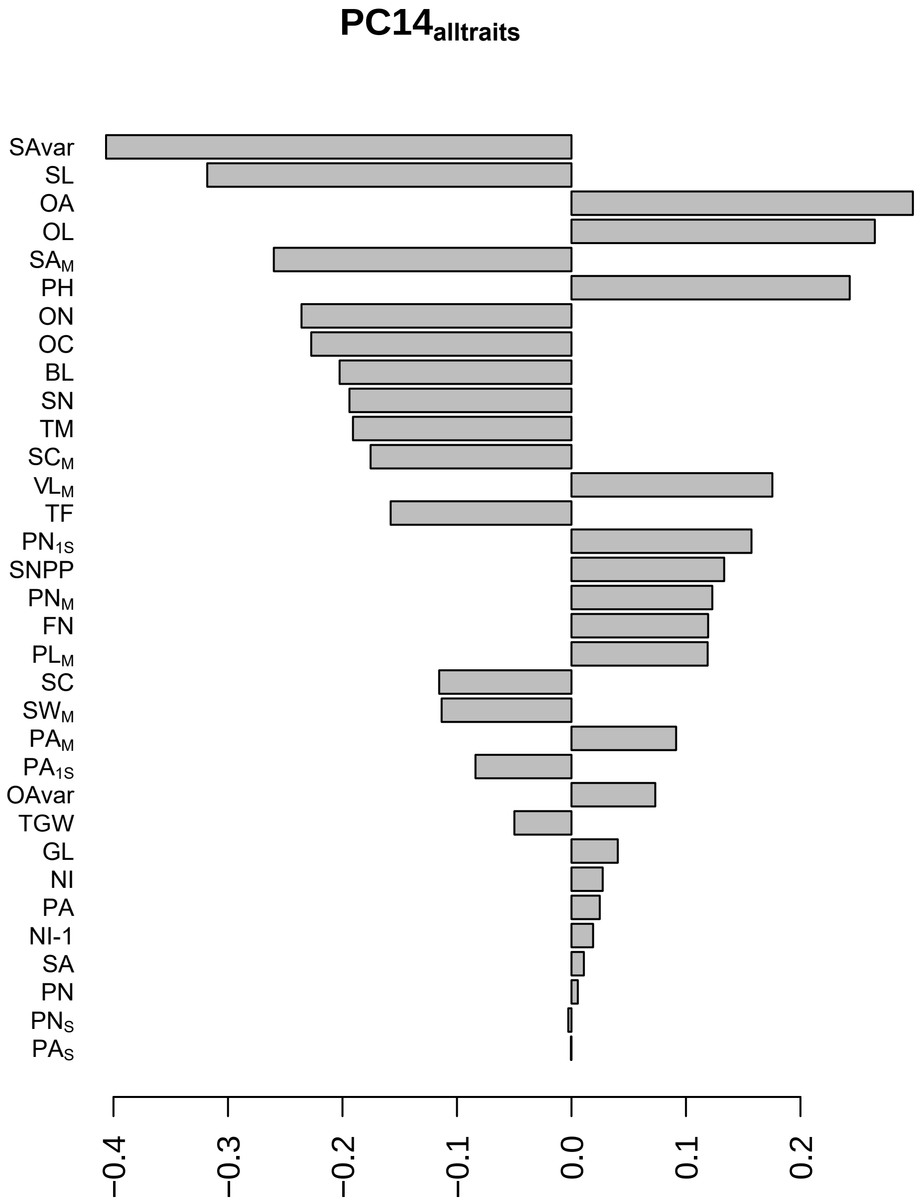
PC14alltraits
SAvar
SL
OA
OL
SAM
PH
ON
OC
BL
SN
TM
SCM
VLM
TF
PN1S
SNPP
PNM
FN
PLM
SC
SWM
PAM
PA1S
OAvar
TGW
GL
NI
PA
NI-1
SA
PN
PNS
PAS
−0.4
−0.3
−0.2
−0.1
0.0
0.1
0.2

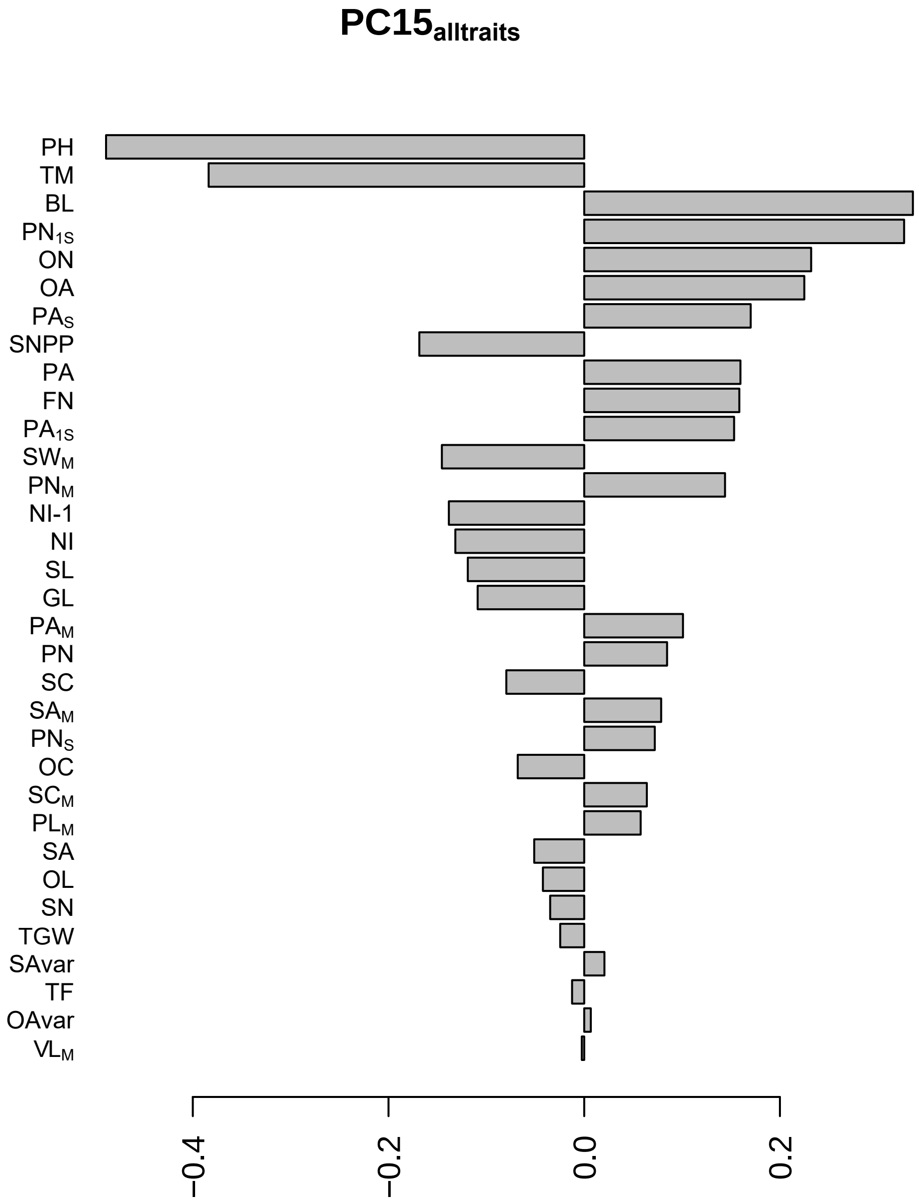
PC15alltraits
PH
TM
BL
PN1S
ON
OA
PAS
SNPP
PA
FN
PA1S
SWM
PNM
NI-1
NI
SL
GL
PAM
PN
SC
SAM
PNS
OC
SCM
PLM
SA
OL
SN
TGW
SAvar
TF
OAvar
VLM
−0.4
−0.2
0.0
0.2
