## Supplemental Table S1 for "Uncovering the ideal plant ideotype for maximising seed yield in *Brassica napus*"

Supplemental Table S1. Ranks the Winter OSR genotypes according to its position within each PC_macro_ (n=5).

| **Genotypes** | **Seed yield (g)** | **PC1_macro_** | **PC5_macro_** | **PC6_macro_** | **PC7_macro_** |
| --- | --- | --- | --- | --- | --- |
| POH 285, Bolko | 21.29 | 18 | 24 | 30 | 34 |
| Canberra x Courage | 20.46 | 4 | 26 | 4 | 21 |
| Norin | 20.42 | 9 | 8 | 32 | 6 |
| Shannon x Winner DH | 20.28 | 25 | 1 | 31 | 8 |
| Verona | 20.21 | 11 | 7 | 7 | 26 |
| Rafal DH1 | 19.98 | 7 | 9 | 1 | 38 |
| Rocket | 19.50 | 3 | 40 | 5 | 3 |
| Madrigal x Recital DH | 19.39 | 19 | 32 | 18 | 4 |
| Capitol | 19.04 | 26 | 12 | 8 | 22 |
| Hansen x Gaspard DH | 18.85 | 13 | 16 | 38 | 10 |
| Inca x Contact | 18.76 | 8 | 29 | 16 | 7 |
| Expert | 18.75 | 5 | 30 | 9 | 17 |
| Huron x Navajo | 18.41 | 1 | 27 | 14 | 13 |
| Matador | 18.32 | 2 | 10 | 2 | 11 |
| Licrown x Express DH | 18.21 | 33 | 28 | 6 | 27 |
| Eurol | 17.80 | 16 | 2 | 22 | 35 |
| Vision | 17.32 | 10 | 22 | 39 | 14 |
| Temple | 17.26 | 12 | 35 | 12 | 2 |
| Dimension | 17.06 | 23 | 19 | 11 | 29 |
| Cabernet | 16.75 | 21 | 39 | 17 | 9 |
| Dippes | 16.64 | 6 | 41 | 20 | 15 |
| Coriander | 16.41 | 22 | 38 | 29 | 28 |
| Apex-93_5 x Ginyou_3 DH | 16.25 | 28 | 20 | 36 | 25 |
| Abukuma Natane | 16.10 | 24 | 4 | 35 | 37 |
| Palmedor | 15.91 | 20 | 21 | 26 | 1 |
| Apex | 15.85 | 38 | 15 | 34 | 12 |
| Lembkes Malchower (Lenora) | 15.26 | 14 | 23 | 23 | 33 |
| Baltia | 15.09 | 31 | 3 | 40 | 32 |
| Cabriolet | 14.98 | 15 | 34 | 15 | 36 |
| Tapidor DH | 14.87 | 35 | 18 | 24 | 20 |
| Kromerska | 14.50 | 34 | 6 | 19 | 18 |
| Slovenska Krajova | 14.46 | 29 | 17 | 25 | 19 |
| Castille | 14.07 | 39 | 5 | 42 | 23 |
| Ramses | 13.70 | 17 | 37 | 28 | 30 |
| Lesira | 13.58 | 36 | 33 | 13 | 40 |
| Excalibur | 13.03 | 40 | 13 | 27 | 5 |
| Quinta | 13.02 | 30 | 25 | 41 | 41 |
| Samourai | 12.51 | 27 | 36 | 3 | 42 |
| Catana | 11.58 | 37 | 42 | 10 | 39 |
| Janetzkis Schlesischer | 11.34 | 32 | 31 | 21 | 31 |
| Bienvenu DH4 | 10.80 | 41 | 14 | 37 | 24 |
| Flash | 5.38 | 42 | 11 | 33 | 16 |
