## Supplemental Table S4 for "Uncovering the ideal plant ideotype for maximising seed yield in *Brassica napus*"

Supplemental Table S4. Ranks the Spring OSR genotypes according to its position within each PC_alltraits_ (n=3).

| **Genotype** | **Seed yield (g)** | **PC1_alltraits_** | **PC6_alltraits_** | **PC7_alltraits_** | **PC10_alltraits_** |
| --- | --- | --- | --- | --- | --- |
| Mazowiecki | 15.99 | 1 | 2 | 21 | 8 |
| Cresor | 15.85 | 4 | 15 | 8 | 17 |
| Tantal | 15.83 | 9 | 1 | 20 | 16 |
| Westar DH | 14.99 | 6 | 14 | 6 | 13 |
| Erglu | 14.82 | 7 | 12 | 3 | 21 |
| Ceska Krajova | 14.70 | 11 | 17 | 9 | 2 |
| N01D-1330 | 14.42 | 3 | 10 | 13 | 9 |
| Topas | 13.90 | 14 | 6 | 10 | 7 |
| Willi | 13.88 | 12 | 13 | 17 | 4 |
| Duplo | 13.48 | 8 | 19 | 5 | 6 |
| Bronowski | 13.43 | 5 | 21 | 1 | 15 |
| Tribune | 13.10 | 10 | 7 | 2 | 19 |
| Drakkar | 12.27 | 15 | 3 | 14 | 18 |
| Helios | 10.34 | 13 | 11 | 12 | 12 |
| Karat | 9.80 | 2 | 4 | 22 | 22 |
| Monty-028 DH | 9.59 | 16 | 22 | 11 | 20 |
| N02D-1952 | 9.23 | 18 | 8 | 16 | 5 |
| Karoo-057 DH | 8.82 | 19 | 20 | 19 | 10 |
| Surpass400-024 DH | 7.67 | 17 | 5 | 7 | 3 |
| Weihenstephaner | 6.36 | 20 | 9 | 15 | 14 |
| Stellar DH | 4.36 | 21 | 16 | 18 | 11 |
| Cubs Root | 3.79 | 22 | 18 | 4 | 1 |
