## Supplemental Table S5 for "Uncovering the ideal plant ideotype for maximising seed yield in *Brassica napus*"

Supplemental Table S5. Percentage of seed yield variation explained by different Partial Least Square (PLS) components for macrotraits for WOSR and SOSR.

| Winter OSR | | Spring OSR | |
| --- | --- | --- | --- |
| PLS components | **Seed yield variation (%)** | **PLS components** | **Seed yield variation (%)** |
| 1 | 44.2 | 1 | 74.0 |
| 2 | 22.1 | 2 | 9.7 |
| 3 | 13.2 | 3 | 5.9 |
| 4 | 8.0 | 4 | 2.0 |
| 5 | 3.0 | 5 | 1.9 |
| 6 | 3.0 | 6 | 1.1 |
| 7 | 1.3 | 7 | 1.2 |
| 8 | 0.8 | 8 | 0.9 |
| 9 | 0.7 | 9 | 0.6 |
