## Supplemental Table S6 for "Uncovering the ideal plant ideotype for maximising seed yield in *Brassica napus*"

Supplemental Table S6: Percentage of seed yield variation explained by different Partial Least Square (PLS) components for alltraits for WOSR and SOSR.

| Winter OSR | | Spring OSR | |
| --- | --- | --- | --- |
| PLS components | **Seed yield variation (%)** | **PLS components** | **Seed yield variation (%)** |
| 1 | 46.3 | 1 | 74.8 |
| 2 | 21.8 | 2 | 8.7 |
| 3 | 13.3 | 3 | 5.5 |
| 4 | 6.1 | 4 | 3.3 |
| 5 | 2.9 | 5 | 1.4 |
| 6 | 2.1 | 6 | 1.2 |
| 7 | 1.8 | 7 | 1.2 |
| 8 | 1.1 | 8 | 0.7 |
| 9 | 0.6 |  |  |
| 10 | 0.6 |  |  |
| 11 | 0.4 |  |  |
