## Supplemental Table S9 for "Uncovering the ideal plant ideotype for maximising seed yield in *Brassica napus*"

Supplemental Table S9. List of macrotrait (n=5) and microtrait (n=3) names and abbreviations measured in the diversity set population.

| Macrotraits | | Microtraits | |
| --- | --- | --- | --- |
| Trait name | **Abbreviation** | **Trait name** | **Abbreviation** |
| Plant height (cm) | PH | Ovule number | ON |
| Number of flowering inflorescences | NI | Ovule area (mm^2^) | OA |
| Number of secondary inflorescences | NI-1 | Ovary length (mm) | OL |
| Time to flowering (days) | TF | Gynoecia length (mm) | GL |
| Number of flowers on the whole plant | FN | Style length (mm) | SL |
| Number of pods on the main inflorescence | PN_M_ | Beak length (cm) | BL |
| Number of pods on a secondary inflorescence | PN_1s_ | Ovule area coefficient of variation (%) | OAcvar |
| Number of pods on secondary inflorescences | PN_s_ |  |  |
| Number of pods on the whole plant | PN |  |  |
| Pod abortion on the main inflorescence (%) | PA_M_ |  |  |
| Pod abortion on a secondary inflorescence (%) | PA_1s_ |  |  |
| Pod abortion in secondary inflorescences (%) | PA_s_ |  |  |
| Pod abortion in the whole plant (%) | PA |  |  |
| Time to maturity (days) | TM |  |  |
| Pod length from 10 pods from the main inflorescence (cm) | PL_M_ |  |  |
| Valve length from 10 pods from the main inflorescence (cm) | VL_M_ |  |  |
| Seed number/ pod from 10 pods from the main inflorescence | SNPP_M_ |  |  |
| Seed area from 10 pods from the main inflorescence (mm^2^) | SA_M_ |  |  |
| Seed compactness from 10 pods from the main inflorescence | SC_M_ |  |  |
| Seed weight from 10 pods from the main inflorescence (g) | SW_M_ |  |  |
| Seed area from the whole plant (mm^2^) | SA |  |  |
| Seed compactness from the whole plant | SC |  |  |
| Seed area coefficient of variation from whole plant (%) | SAcvar |  |  |
| Thousand grain weight (g) | TGW |  |  |
| Estimated total seed number from the whole plant (by TGW) | SN |  |  |
| Seed oil content from the whole plant (%) | OC |  |  |
| Seed weight from the whole plant (seed yield, g) | SY |  |  |
