## Supplemental Table S10 for "Uncovering the ideal plant ideotype for maximising seed yield in *Brassica napus*"

| Trait | Transformation |
| --- | --- |
| Plant height | square root |
| Number of flowering inflorescences | natural log |
| Number of secondary inflorescences | natural log |
| Number of flowers on the whole plant | natural log |
| Number of pods on secondary inflorescences | square root |
| Number of pods on the whole plant | square root |
| Pod abortion on the main inflorescence | logit with offset |
| Pod abortion in secondary inflorescences | logit with offset |
| Pod abortion in the whole plant | logit with offset |
| Ovary length | natural log |
| Time to maturity | natural log |
| Seed area from 10 pods from the main inflorescence | natural log |
| Seed compactness from 10 pods from the main inflorescence | logit |
| Seed area from the whole plant | natural log |
| Seed compactness from the whole plant | logit |
| Estimated total seed number from the whole plant (by TGW) | natural log |
| Seed oil content from the whole plant | logit, with reference of 50 |

Supplemental Table S10. List of transformations applied in order to satisfy homogeneity of variances.
